# Liver lipophagy ameliorates nonalcoholic steatohepatitis through lysosomal lipid exocytosis

**DOI:** 10.1101/2022.02.22.481456

**Authors:** Yoshito Minami, Atsushi Hoshino, Yusuke Higuchi, Masahide Hamaguchi, Yusaku Kaneko, Yuhei Kirita, Shunta Taminishi, Akiyuki Taruno, Michiaki Fukui, Zoltan Arany, Satoaki Matoba

## Abstract

Nonalcoholic steatohepatitis (NASH) is a progressive disorder with aberrant lipid accumulation and subsequent inflammatory and profibrotic response. Therapeutic efforts at lipid reduction via increasing cytoplasmic lipolysis unfortunately worsens hepatitis due to toxicity of liberated fatty acid. An alternative approach could be lipid reduction through autophagic disposal, i.e., lipophagy. We engineered a synthetic adaptor protein to induce lipophagy, combining a lipid droplet-targeting signal with optimized LC3-interacting domain. Activating hepatocyte lipophagy in vivo strongly mitigated both steatosis and hepatitis in a diet-induced mouse NASH model. Mechanistically, activated lipophagy promoted the excretion of lipid from hepatocytes via lysosomal exocytosis, thereby suppressing harmful intracellular accumulation of nonesterified fatty acid. A high-content compound screen identified alpelisib and digoxin, clinically-approved compounds, as effective activators of lipophagy. Administration of alpelisib or digoxin in vivo strongly inhibited the transition to steatohepatitis. These data thus identify lipophagy as a promising therapeutic approach to prevent NASH progression.

## Introduction

Lifestyle changes have led to worldwide increases in nonalcoholic fatty liver disease (NAFLD), with a prevalence now estimated to be around 25%^1^. Twenty to thirty percent of patients with NAFLD progress to nonalcoholic steatohepatitis (NASH), which in turn can progress to liver cirrhosis liver and eventually hepatocellular carcinoma. NASH has become the leading cause of liver failure in the US ^2^. The pathogenesis of NASH development is complicated, involving multiple hits ^3^. Excessive accumulation of lipid and abnormal fatty acid metabolism generates reactive oxygen species and endoplasmic reticulum (ER) stress. Infiltration by inflammatory cells and the subsequent fibrotic response drive NASH progression and contribute to increased mortality ^4^. Circulating inflammatory monocytes are recruited via chemokine receptors^5^ and differentiate into liver macrophages, followed by activation of hepatic stellate cells (HSCs) and accumulation of extracellular matrix in the liver^6^. Despite the high prevalence and clinical importance of NAFLD, there is no approved drug for the treatment of NAFLD/NASH ^7,8.^

Aberrant accumulation of lipid droplets (LDs) is the initial step of NAFLD/NASH pathogenesis, and it is thought that NASH is almost never resolved without improvement in steatosis ^9^. However, dysregulated breakdown of LDs, despite lowering LD content, can contribute to disease progression. For example, studies with ablation of Perilipin5, which normally prevents adipose triglyceride lipase (ATGL) transposition to LDs and thus prevents lipolysis^10^, show that activated lipolysis reduces hepatosteatosis but worsens hepatic lipotoxicity and insulin resistance ^11,12^. Mechanistically, it is thought that the altered fatty acid composition caused by inefficient incorporation of fatty acids into LD-containing triacylglycerol drives NAFLD ^13^.

Lysosomes provide an alternative pathway to degrade LDs, as well as other intracellular organelles and molecules, via various lysosomal hydrolases such as lipase, proteases, glycosidases, and nucleotidases ^14^. Lysosomal acid lipase, LIPA (also known as LAL) degrades lysosomal lipids, and mutations in LIPA cause accumulation of LDs in various organs ^15,16,^ demonstrating the importance of lipophagy for lipid homeostasis. Lipophagy was first demonstrated in hepatocytes under nutrient deprivation^17^, but accumulating evidence shows that lipophagy contributes to LD degradation in numerous cell types, and that aberrant lipophagy is common in numerous diseases, including fatty liver, obesity and cancers ^18^. However, the molecular mechanism of how autophagy targets LDs remains poorly understood, with most mechanistic insight extrapolated from studies of bulk autophagy. In the liver, models of hepatocyte-specific autophagy deficiency have yielded complicated results, in part due to the lipogenic aspect of autophagosomes. For example, deletion of Atg7 or FIP200 impaired de novo lipogenic program by liver X receptor and protected from high-fat diet-induced fat accumulation ^4,19–22.^ In contrast, activation of autophagy via overexpression of the transcription factor EB (TFEB) or by ablation of Rubicon clearly demonstrated the protective role of liver autophagy on NASH ^23,24^. However, the specific role of lipophagy in these studies was not fully explored.

To definitely and specifically evaluate the effect of LDs disposal by autophagy, we developed here synthetic adaptor proteins, fusing LD-targeting domains with optimized LC3 interacting regions (LIR), to induce selective lipophagy of LDs and evaluate its impact on liver lipid homeostasis. Our adaptor protein efficiently activated lipophagy and protected against NFALD and NASH in mice fed high fat with low methionine low choline diet (MCD). Mechanistically, upregulated lipophagy was coupled with lysosomal exocytosis, rather than fatty acid catabolism, to ultimately reduce liver fatty acid content and consequential inflammation and fibrosis. Finally, high throughput compound screening for lipophagy activators identified FDA-approved drugs that provide proof-of-concept for the potential clinical application of our findings.

## Results

### Development of an optimized lipophagy adaptor protein

We engineered a lipophagy adaptor protein, designed to induce selective autophagy of lipid droplets, by fusing a lipid droplet-targeting signal (LDTS) with an autophagosome recruiting domain, i.e., an engineered LC3-interacting region (eLIR) (Fig. 1a). To first identify the LIR that best recruits autophagosome, LIR-containing proteins were fused with a mitochondrial outermembrane-targeting signal, and the level of mitophagy was analyzed with flow cytometry and the pH-indicator, mKeima ^25^, which loses green fluorescence when in the presence of lysosomal low pH. Among sequestosome-1-like receptors, optineurin (OPTN) exhibited the highest power to recruit autophagosome when expressed in the surface of mitochondria, and seemed more efficient than the previously reported AMBRA1-based mitophagy adaptor protein ^26^ (Supplementary Fig. 1a). OPTN is multi-functional protein modulating a number of signaling pathways like NF-κB, as well as trafficking of vesicles and proteins ^27^. To prevent unintended effects of expressing intact OPTN, non-essential sequences were excluded, and residues 120-190 were identified as the minimal sequence necessary to promote autophagic degradation (Supplementary Fig. 1b). TBK1-mediated serine phosphorylation near the OPTN LIR is known to induce conformational alteration that enhances LC3 binding affinity ^28^. In addition to phosphomimetic mutation in 5 residues, replacement of phenylalanine with tryptophan in the LIR of OPTN also enhances the affinity to LC3 by increasing the binding enthalpy^29^. We therefore adopted both of these modifications, and in addition, modified the adaptor protein as a free form using SunTag system^30^. These modifications promoted mitophagy significantly more efficiently than did wild-type OPTN aa120-190 (Supplementary Fig. 1c). We therefore adopted this configuration as our engineered LIR (eLIR).

**Figure 1.**
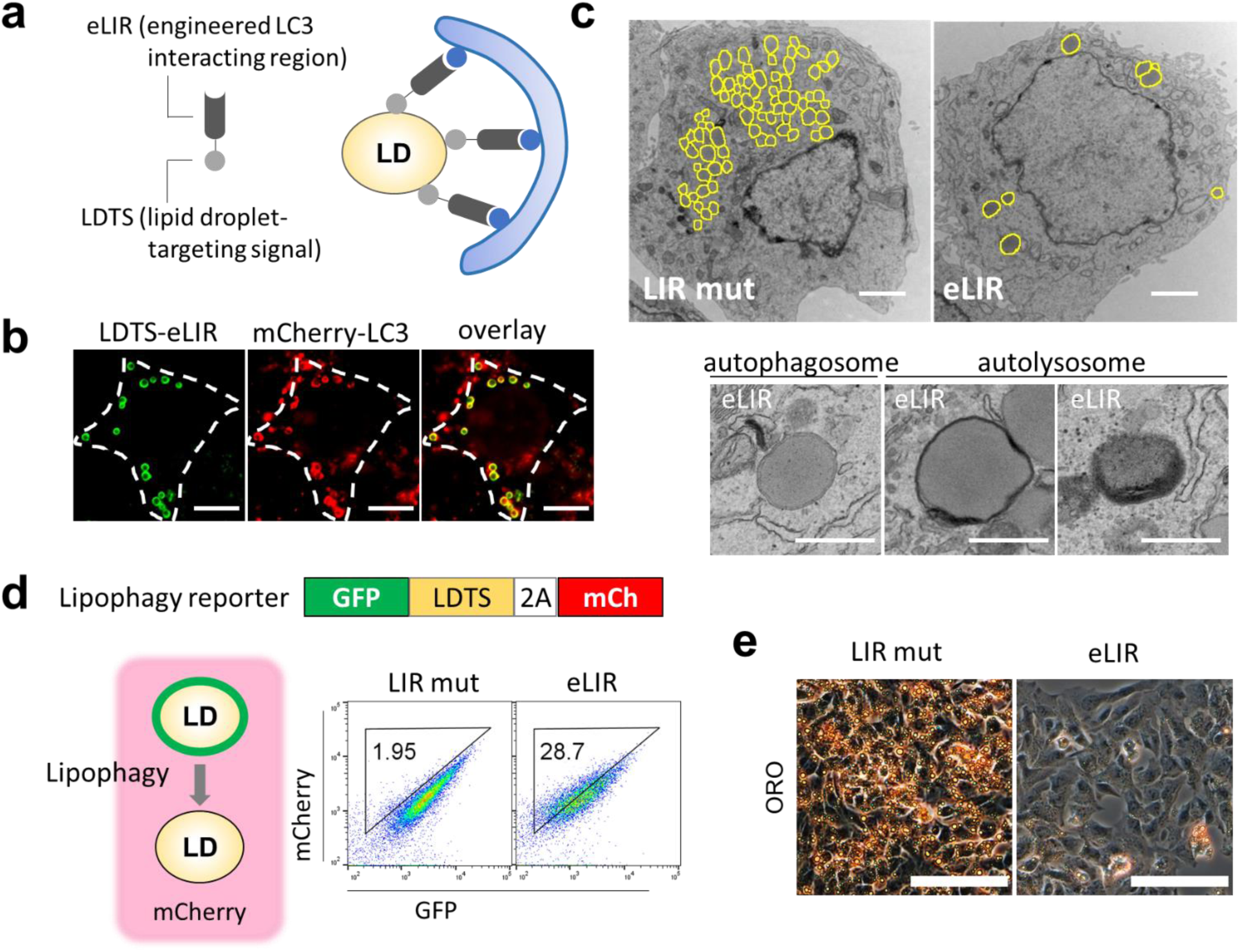
Synthetic adaptor protein to induce lipophagy. (**a**) Schema of lipophagy-inducing synthetic adaptor protein. The lipid droplet-targeting sequence (LDTS) from PLIN1 was fused with autophagosome recruiting engineered LC3 interacting region (eLIR). (**b**) 3T3L1 adipocytes expressing LDTS-eLIR-GFP and mCherry-LC3 were analyzed by fluorescence microscopy. Scale bars, 10 µm. (**c**) 3T3L1 adipocytes expressing LDTS-mutant LIR (LIR mut) or LDTS-eLIR were analyzed by electron microscopy. Lipid droplet is highlighted in yellow. Scale bars, 2 µm and 500 nm. (**d**) Lipophagy was quantified by flow cytometry for LDTS-GFP and internal control of mCherry in HepG2 cells expressing LDTS-LIR mut or LDTS-eLIR treated with oleic acid (OA) for 24 h. (**e**) HepG2 cells expressing LDTS-LIR mut or LDTS-eLIR were treated with OA for 24 h, and lipid droplets were stained with oil red O. Scale bars, 100 µm

Next, we optimized the lipid droplet (LD) targeting signal (LDTS), using flow cytometry-based lipophagy assays in 3T3-L1 adipocytes. mKeima was fused with the LD protein perilipin1 to localize mKeima to LDs. Lipophagy was clearly induced in constructs fusing the eLIR with perilipin1, but little with LDTS from hepatitis C virus^31^ or PNPLA5 (Supplementary Fig. 2a) ^32^. The core domain of perilipin1, which contains the LDTS ^33^, was sufficient to induce lipophagy as efficiently as the full length construct, and was more efficient than the LDTS from perilipin 5 (Supplementary Fig. 2b). Fluorescent imaging confirmed that the LDTS-eLIR was present on the surface of LDs and colocalized with LC3 labelled with mCherry (Fig. 1b), indicating targeting of LDs to the autophagosome. Electron micrographs exhibited LDs in autophagosome and autolysosome and the number of LDs was decreased in adipocytes expressing LDTS-eLIR (Fig. 1c). Lipophagy was also evaluated with administration of oleic acid to lipophagy-reporter HepG2 hepatocytes that express GFP-perilipin1 and mCherry as a lipophagy reporter: when lipophagy occurs, the LD-associated GFP loses fluorescence in lysosomal acid conditions, while the mCherry signal is retained. Expression of the LDTS-eLIR construct strikingly activated lipophagy in these cells (Fig. 1d) and reduced the accumulation of LDs, as seen by staining with oil red O (Fig. 1e) or BODIPY (Supplementary Fig. 3a), or by flow cytometry (Supplementary Fig. 3b). Our LDTS-eLIR construct activated lipophagy more reliably and selectively than did TFEB overexpression ^23^ or starvation (Supplementary Fig. 3c).

### Activation of hepatocyte lipophagy reverses steatohepatitis

To test the impact of activating lipophagy on NAFLD/NASH pathology, the LDTS-eLIR construct was engineered into serotype 8 adeno-associated virus (AAV8) with liver specific thyroxine-binding globulin promoter (pTBG). AAV8s carrying LDTS-eLIR and LDTS-mutant LIR (LIR mut) were injected in mice after 6-weeks of feeding high fat with low methionine low choline diet (MCD) as a NASH model ^34^. Mice were then switched to normal chow, and analyzed 2-week later (Fig. 2a). The introduction of LDTS-eLIR did not affect food intake or body weight, but it did reduce liver weight significantly (Fig. 2b-d). This was accompanied by markedly decreased liver steatosis, as evidenced by H&E histology, oil red O staining, and quantification of triglyceride and nonesterified fatty acid content (Fig. 2e,f). Importantly, serum lipid profile remained unaltered in animals treated with LDTS-eLIR (Fig. 2g). The reduced steatosis in the liver was associated with evidence of mitigated liver injury (Fig. 2h). Gene set enrichment analysis of RNAseq of liver tissue indicated that activation of lipophagy mitigated profibrotic and proinflammatory responses (Fig. 2i,j). Consistent with this, α-smooth muscle actin (α-SMA) protein expression, a marker of HSC activation, and fibrosis staining (Sirius red) were strongly attenuated in the lipophagy-induced liver (Fig. 2k-m). The induction of lipophagy also led to higher expression of mitochondrial oxidative phosphorylation (OXPHOS) genes that might reflect milder NASH pathology (37) (Fig 2j).

**Figure 2.**
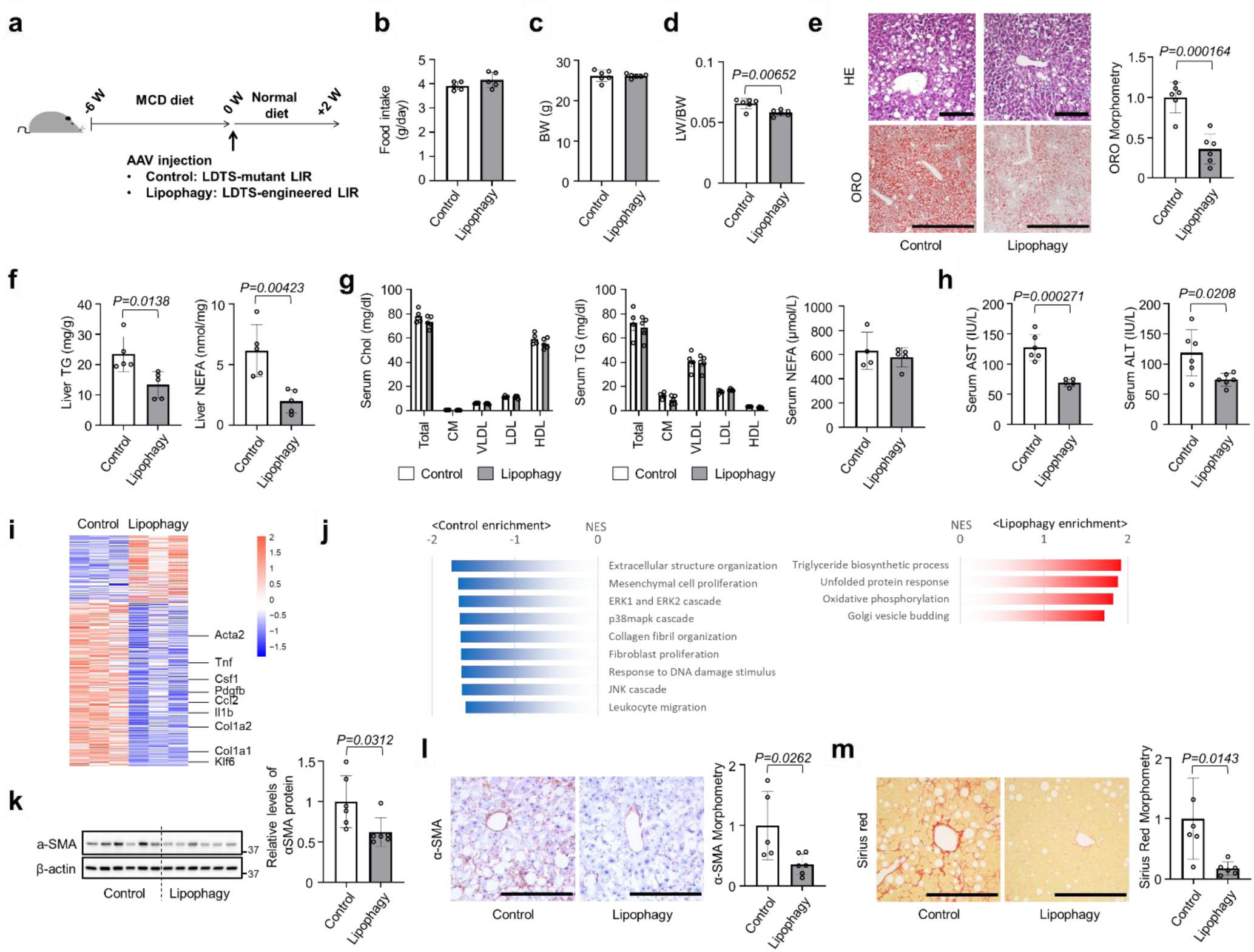
Lipophagy prevented nonalcoholic steatohepatitis in mice fed MCD diet. (**a**) Study protocol of NASH model with high fat with low methionine low choline (MCD) diet and the induction of lipophagy with AAV-mediated gene delivery into hepatocytes. (**b**-**d**) Food intake (b), Body weight (c) and liver/body weight (LW/BW) ratios (d) of lipophagy-induced NASH model mice. (**e**) Hematoxylin/eosin or oil red O staining of liver sections. Scale bars, 1 mm. (**f**) Liver TG and NEFA content. (**g**) Serum cholesterol and TG in 4 major fractions and NEFA content. (**h**) Serum AST and ALT levels. (**I**,**j**) RNA sequence and gene set enrichment analysis of liver tissues. (**k**) Immunoblots assessing the expression of αSMA in liver tissues. (**l**,**m**) Immunohistochemistry of αSMA (l) and sirius red staining (m) of liver sections and morphometry analysis of signal positive area. Scale bars, 200 µm. All data are presented as mean±SD (n=6 per group). *P* values calculated by two-sided unpaired *t*-test.

To test the prediction that the protection from NAFLD conferred by LDTS-eLIR requires the autophagy machinery, the hepatoprotective effect of LDTS-eLIR was assessed in the autophagy-deficient settings of liver-specific ATG5 knockout mice or chloroquine (CQ)-treated mice. ATG5 gene deletion was achieved by concurrent gene delivery of Cre recombinase with the LDTS-LIRs by the AAV8 pTBG system (Supplementary Fig. 4a,b). In the absence of liver ATG5, the expression of LDTS-eLIR protein induced little alteration of LDs, liver triglycerides, and nonesterified fatty acids (Supplementary Fig. 4c,d). CQ is a classic inhibitor of autophagy by disturbing autophagosome fusion with lysosomes^35^. Like deletion of ATG5, treatment with CQ also prevented the lipid-lowering effect of LDTS-eLIR in HepG2 cells (Supplementary Fig. 4e) and livers from mice fed with MCD diet (Supplementary Fig. 4f-h). These results thus indicate that both autophagosome formation and subsequent fusion with lysosome are essential for the hepatoprotective effect of LDTS-eLIR.

### Activation of lipophagy promotes lysosomal exocytosis of lipids

To investigate the fate of lipids removed by lipophagy, we first assessed lipid catabolism in livers with LDTS-eLIR. Lipophagy-induced liver tissue showed no overall significant alteration of transcription of fatty acid oxidation (FAO), lipolysis, or lipid biosynthesis genes, despite the higher expression of mitochondrial respiration-related genes (Supplementary Fig. 5a,b). Functionally, there was no difference in FAO enzymatic activity of liver lysates, nor in the serum level of β-OHB, a marker of excess fatty acid oxidation in the liver (Fig. 3a,b). FAO activity in cultured HepG2 cells, analyzed using a quenched fluorescent probe that is activated by the sequential enzyme reactions of FAO^36^, also revealed no changes in FAO in lipophagy-activated cells (Supplementary Fig. 5c). Oxidation thus seems unlikely to be the ultimate fate of lipids removed by lipophagy.

**Figure 3.**
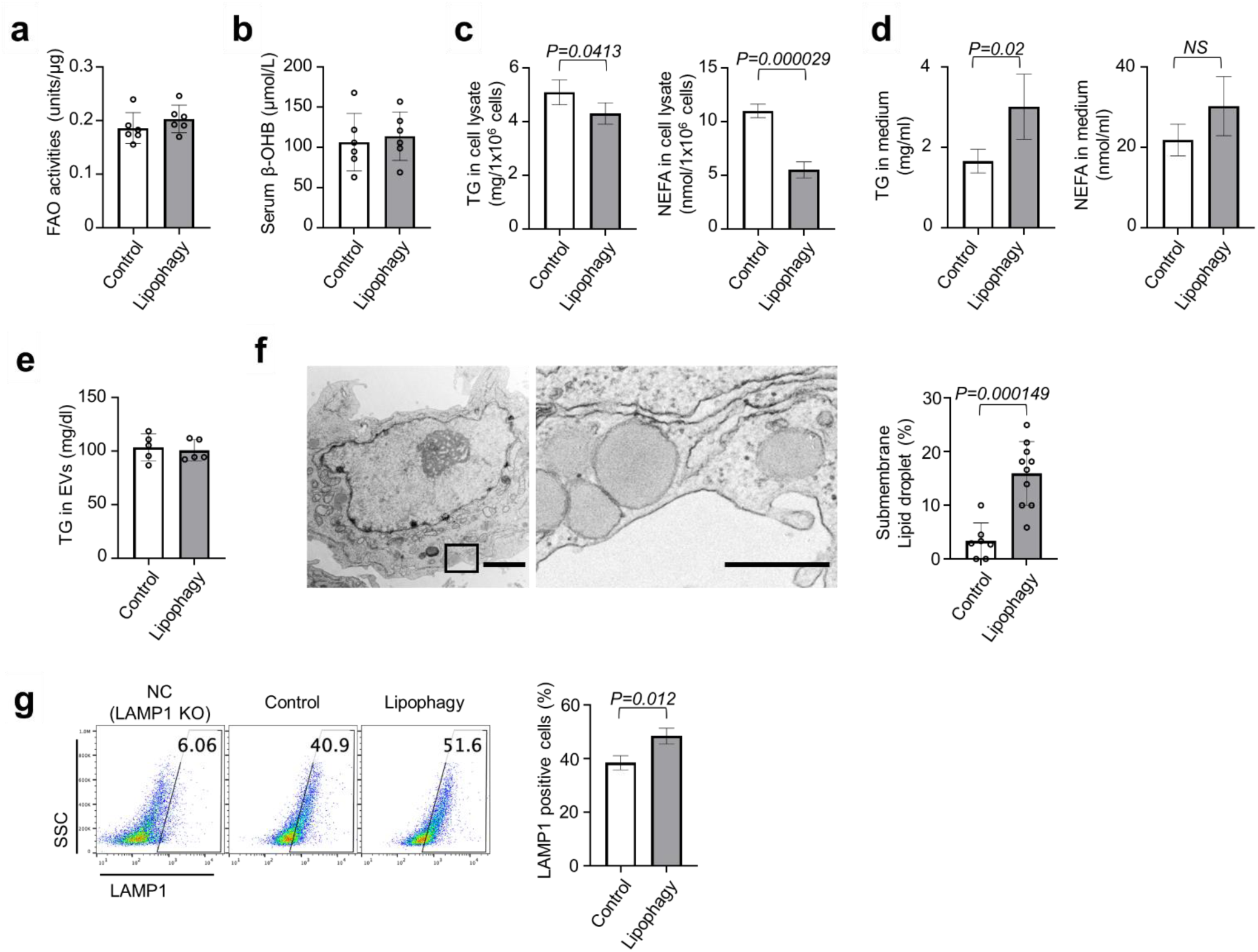
Lipophagy excreted LDs through the lysosomal exocytosis. **(a)** Fatty acid oxidation (FAO) activities of liver tissues in mice fed MCD diet for the NASH model. **(b)** The levels of serum ß-OHB in mice fed MCD diet for the NASH model (a,b; n=6 per group). (**c**,**d**) HepG2 cells expressing LDTD-LIR mut (control) or eLIR (lipophagy) were cultured with FFAs (OA+PA) containing medium for 12 h and then incubated in FBS-free medium for 6 h. TG and NEFA of cell lysate (c) and culture supernatant (d) were measured (n=4 per group). (**e**) TG content in serum exosomes in a mouse model of NASH fed with MCD diet (n=5 per group) (**f**) Electron micrographs of lipophagy-induced hepatocyte. Right image is magnified view of left black square. LDs localized close to cellular membrane was quantified. Scale bars, 2 µm and 500 nm (n=10 cells per group). (**g**) Flow cytometry for Lamp1 in the surface of HepG2 cells expressing LDTD-LIR mut (control) or eLIR (lipophagy). LAMP1 KO cells were used as a negative control (n=3 per group). All data are presented as mean±SD. *P* values calculated by two-sided unpaired *t*-test. Abbreviations: ß-OHB; ß-Hydroxybutyric Acid.

While cellular triglyceride and nonesterified fatty acid were decreased upon induction of lipophagy in oleic acid-loaded HepG2 cells (Fig. 3c), levels of triglyceride in the medium were increased (Fig. 3d). We thus hypothesized that activation of lipophagy caused the induction of lipid export, thereby contributing to liver protection. In mice fed with MCD diet, lipophagy did not change the amount of triglyceride in extracellular vesicles (EVs) (Fig. 3e), suggesting alternative pathways of secretion. Electron micrograph of lipophagy-induced hepatocytes showed that LDs were localized close to the cell membrane (Fig. 3f). In light of these observations, and the essentiality of autophagosome fusion with lysosomes in the protective effect of LDTS-eLIR in CQ-treated mice experiments (Supplementary Fig. 4f-h), we focused our attention on lysosomal exocytosis.

Lysosomal exocytosis causes the accumulation of LAMP1 at the cell surface ^37^, and introduction of LDTS-eLIR in HepG2 cells significantly increased the amount of cell surface LAMP1 (Fig. 3g). Interestingly, this evidence of lysosomal exocytosis was not observed after activation of other forms of selective autophagy, such as mitophagy and pexophagy (the latter induced by fusion protein of the localization motif of the peroxin Pex13 ^38^ and eLIR, and monitored by peroxisome-localized mKeima with the carboxyl-terminal amino acid sequence serine-lysine-leucine (SKL) ^39^ (Supplementary Fig. 6a,b). To better understand the mechanism underlying the activation of lysosomal exocytosis by lipophagy, we focused on the Ca2+ signal-dependent lysosomal fusion with cell membrane ^40,41^. Knockout of the lysosome Ca2+ channel, TRPML1, or the cell membrane Ca2+ sensor, SYT7, completely blocked the induction of cell surface LAMP1 by the activation of lipophagy (Fig. 4a,b). Similarly, lipophagy-mediated excretion of triglyceride was completely blocked in SYT7 or TRPML1 knockout cells (Fig. 4c). We conclude that activation of lipophagy leads to Ca2+-dependent lysosomal exocytosis of triglycerides.

**Figure 4.**
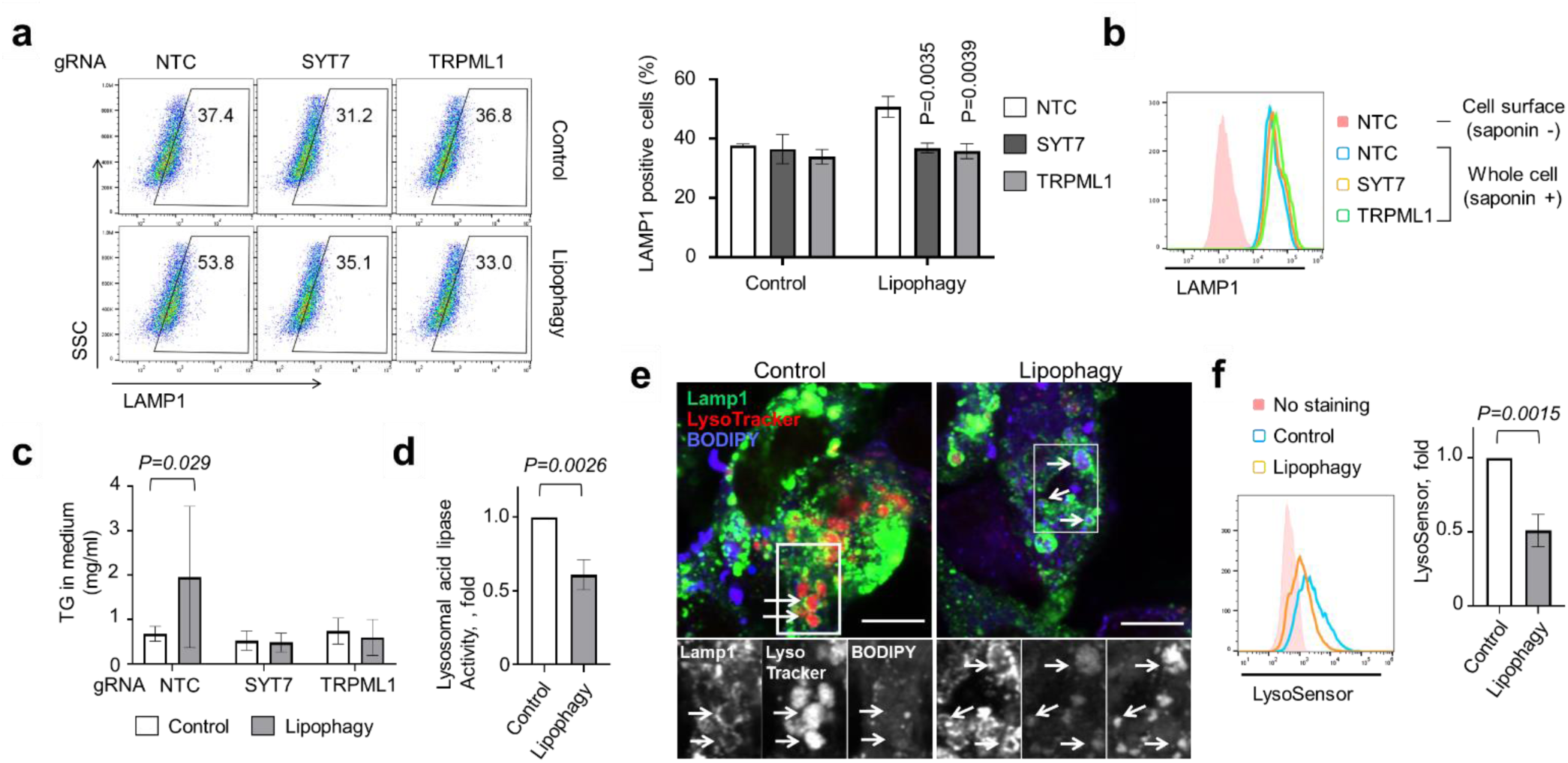
Deacidification activated lysosomal exocytosis. (**a**) Flow cytometry for Lamp1 in the surface of control and lipophagy-induced HepG2 cells. SYT7 or TRPML1 was knockout by CRISPR-Cas9 (n=3 per group). (**b**) Flow cytometry to evaluate whole cell expression level of LAMP1 in saponin-treated cells. (**c**) TG in the medium of control and lipophagy-induced HepG2 cells. Cells were cultured with FFAs (OA+PA) containing medium for 12 h and then maintained in FBS-free medium for 6 h (n=3-9 per group). (**d**) Lysosome acid lipase activity in control and lipophagy-induced HepG2 cells (n=3 per group). (**e**) Lamp1, LysoTracker Red DND-99 and BODIPY co-staining in control and lipophagy-induced HepG2 cells. Scale bar, 10 mm. (**f**) Flow cytometry for LysoSensor Green DND-189 staining of control and lipophagy-induced HepG2 cells (n=3 per group). All data are presented as mean±SD. *P* values calculated by two-sided unpaired *t*-test.

Inhibition of mTORC1 is one of the signals for TRPML1-mediated Ca2+ efflux ^42^, but no alteration of mTORC1 activity was observed after activation of lipophagy (Supplementary Fig. 6c). Next, we examined the contribution of fatty acid to Ca2+ signal in cells expressing cell surface-localized mutant TRPML1 channels ^43^. Monitoring Fura-2 revealed that neither oleic acid nor fatty acid mixture stimulated TRPML1 to increase cytosolic Ca2+ (Supplementary Fig. 6d). In considering other possible triggers for lysosomal exocytosis, we noted that, in the assay for mKeima-based lipophagy, the shift of acidic signal was small as compared with the case of mitophagy (Supplementary Fig. 1 and 2). The GFP-based assay also exhibited relatively smaller signal alteration, and additional starvation, known to activate lysosomal acidification^44^, increased the signal (Supplementary Fig. 3c). These observations suggested a defect in lysosomal acidification upon induction of lipophagy. Lysosomal enzymes function in acidic pH, and even small increases in pH impair enzyme activity^45^. Consistent with this, lysosomal lipase activity, measured with self-quenched lipase substrates ^46^, was reduced in lipophagy-induced cells (Fig. 4d). Staining with a cell-permeable and acidotropic LysoTracker Red DND-99 showed that lipid-containing lysosome had decreased signal in lipophagy-induced cells (Fig. 4e), and lipophagy reduced fluorescence by LysoSensor, a dye sensitive to pH alteration (Fig. 4f). These results thus indicate that lipophagy disturbs lysosomal acidification, likely consequently stimulating Ca2+ release through TRP channel ^47^ and subsequent lysosomal exocytosis.

Finally, we noted that if activation of liver lipophagy promotes secretion of lipids, a concern can be raised of ectopic lipid accumulation and lipotoxicity. However, we observed no hypertrophy of white adipose tissue and no lipid accumulation in skeletal muscle and kidney (Supplementary Fig. 7a-e). Similarly, glucose and insulin profiles showed no significant differences upon activation of liver lipophagy in animals subjected to a MCD diet (Supplementary Fig. 7f-i).

### High-throughput compound screen identified lipophagy activators that suppress NASH progression

We performed a small-molecule screen to identify activators of lipophagy, with an eye to developing lipophagy-based NASH therapy. HepG2 hepatocytes stably expressing GFP-mCherry-LDTS were generated for image-based screening of lipophagy (Fig. 5a). Normal LDs are observed as GFP and mCherry double positive dots, whereas LDs undergoing lipophagy lose GFP signal and become mCherry single positive dots. As positive control, transduction of LDTS-eLIR strikingly increased lipophagy in this assay (Fig. 5b). In the imaging analyses, lipophagy score was defined as a subtraction of total fluorescence amount of GFP from that of mCherry in mCherry positive area per cell. LDTS-eLIR expression exhibited about 50 folds-increase in lipophagy score in this analysis (Fig. 5c), consistent with the result of flow cytometry (Fig. 5b). We next screened approximately 3500 small molecules, including 1600 clinically-approved drugs and 1900 validated compounds provided by Drug Discovery Initiative in The University of Tokyo. Among 39 primary hits defined as 3< robust Z score, there were 3 digitalis derivatives, 3 mTOR pathway inhibitors, and 1 PI3K inhibitor (Fig. 5d and Supplementary Table 1). Twenty-three out of 39, digoxin and other PI3K-mTOR inhibitors were validated and also tested for impact on lipid lowering under oleic acid containing culture conditions (Fig. 5e).

**Figure 5.**
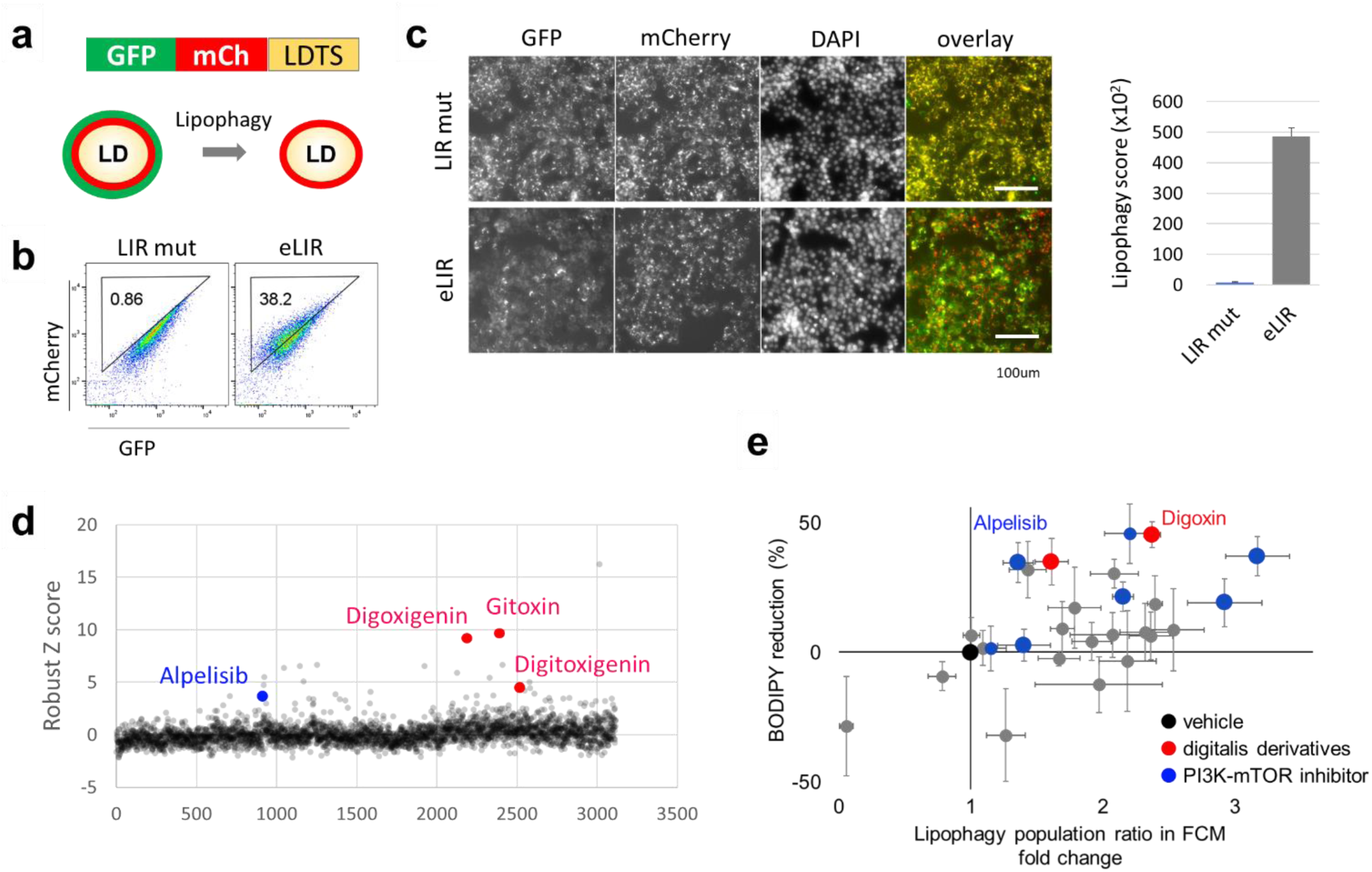
High content compound screening to find lipophagy activators. (**a**) Schematic diagram showing the image-based reporter system to quantify lipophagy activity level. GFP loses its fluorescence in autolysosomes resulting in a decrease in total fluorescent intensity of GFP in mCherry positive area. (**b**) The quality of Lipophagy reporter system was confirmed by flow cytometry with LDTS-LIRs. (**c**) Automated imaging of lipophagy reporter cells expressing LDTS-LIRs as controls. (**d**) Robust Z score of approximately 3500 small molecules library. Toxic compounds defined as cell number < 80% were excluded. (**e**) Individual validation of lipophagy and assessment of intracellular lipid lowering effect among 23 out of 39 primary hits, digoxin and other PI3K-mTOR inhibitors.

We chose two validated hit compounds for further in vivo studies: alpelisib and digoxin. Administered of these clinically-approved compounds to mice fed MCD diet (Fig. 6a) had no impact on body weight but significantly reduced hepatic steatosis, as seen by oil red O staining, and quantified triglyceride and nonesterified fatty acid content (Fig. 6b-d). There were no alterations in serum lipid levels except mild triglyceride increase in alpelisib treatment (Fig. 6e). Alpelisib and digoxin both attenuated inflammatory and profibrotic gene expression, and suppressed liver fibrosis (Fig. 6f-h). Treatment of liver-specific ATG5 knockout mice fed MCD diet with either drug failed to elicit any reduction of liver lipid profiles (Fig. 6i,j), indicating that the protection afforded by alpelisib or digoxin treatment is indeed mediated by lipophagy.

**Figure 6.**
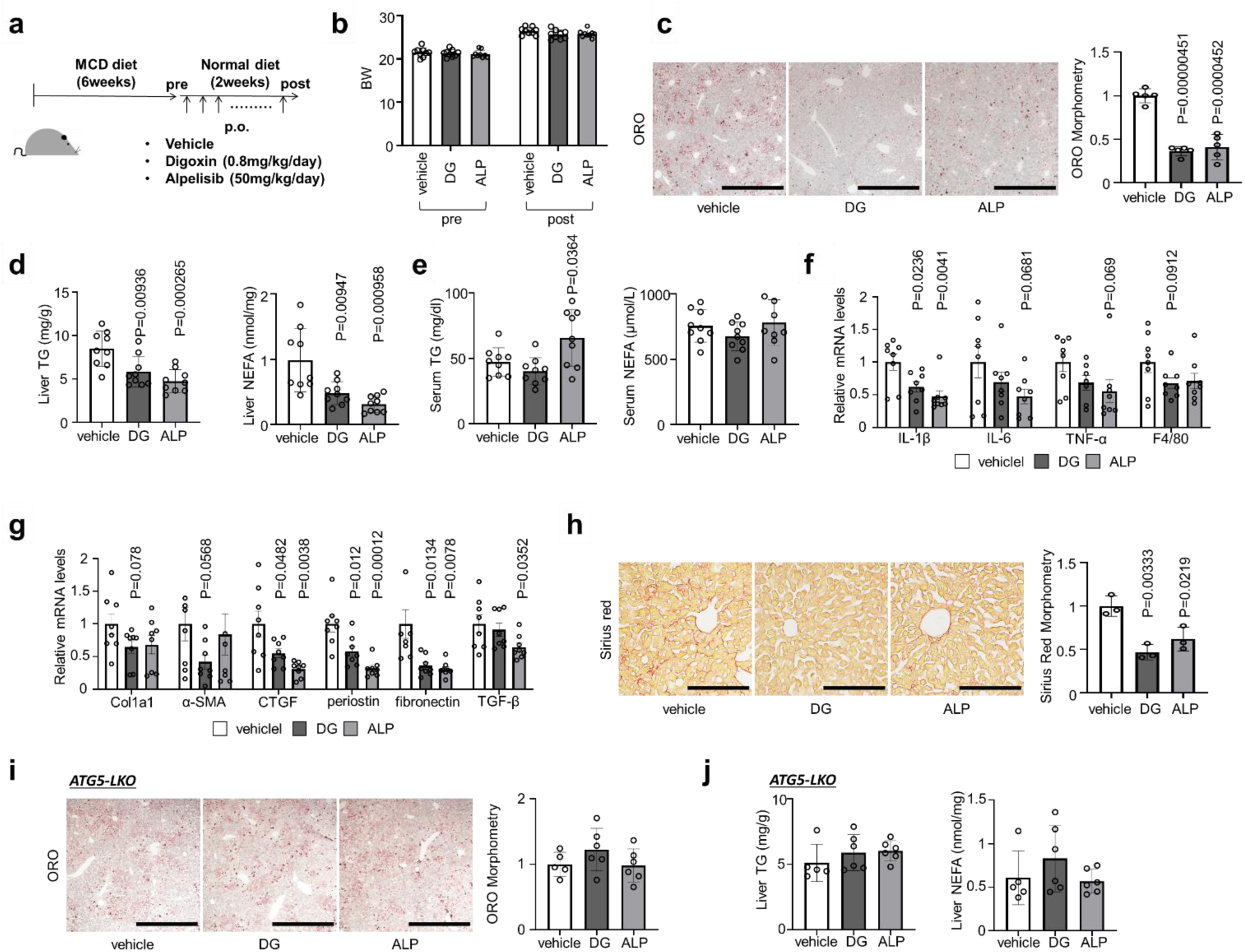
Alpelisib and digoxin prevented the progression of NASH in mice fed MCD diet. (**a**) Study protocol of drug administration in NASH model mice with MCD diet. (**b**) Body weight of control and drug treated mice. (**c**) ORO staining of liver sections. Scale bars, 1 mm. (**d**) Liver TG and NEFA content. (**e**) Serum TG and NEFA levels. (**f**,**g**) Real-time PCR assessing the expression of inflammation (f) and fibrosis (g) related genes in liver tissue. (**h**) Sirius red staining of liver sections. Scale bars, 200µm. (**i**) ORO staining of liver sections of liver-specific ATG5 knockout mice with MCD diet. (**j**) Liver TG and NEFA content of liver-specific ATG5 knockout mice with MCD diet. Data are presented as mean±SD of n=8-9 (a-h) and n=5-6 (i-j). *P* values calculated by two-sided unpaired *t*-test relative to vehicle.

Finally, we retrospectively examined the effect of digoxin administration on indices of fatty liver in human subjects. Digoxin is mainly used in the treatment of heart failure and atrial fibrillation, both of which are associated with NAFLD ^48^. We conducted a nested case-control study, matching for age, gender, BMI, and atrial fibrillation, and excluding patients with heart failure (NYHA class II-IV). A total of 16,172 patients who underwent abdominal ultrasonography in the hospital of Kyoto Prefectural University of Medicine between 2011 and 2020 were included in this study. Among these subjects, 548 were excluded due to missing data or age more than 85 or less than 20 years old, leaving a total of 15,624 patients, of which 282 were actively receiving digoxin treatment. The case-control cohort was selected 1:1, 240 cases in each group. Fatty liver was diagnosed by hepatorenal echo contrast and liver brightness on abdominal ultrasonography ^49^. Strikingly, the prevalence of fatty liver was significantly lower in subjects receiving digoxin than those not receiving digoxin (Table 1).

**Table 1.** Prevalence of fatty liver subjects with Digitalis preparation and those without.

|  | Control | Digoxin | *P* value |
| --- | --- | --- | --- |
| Number of subjects | 240 | 240 |  |
| Fatty liver | 23.8% (57) | 15.4% (37) | 0.025 |
| Atrial fibrillation | 75.0% (180) | 75.0% (180) |  |
| Male | 62.5% (150) | 62.5% (150) |  |
| Age, years | 71.3±9.6 | 71.3±9.6 | 0.96 |
| BMI, kg/m<sup>2</sup> | 21.3±3.1 | 21.3±3.1 | 0.88 |

Continuous variables are expressed as mean± SD and are compared by *t*-test. Categorical variables are expressed as % and number, and are compared by Mantel-Haenzel test.

## Discussions

We show here that liver LDs can be safely and efficiently eliminated by lipophagy, which processes lipids to lysosomal exocytosis rather than to catabolism by mitochondrial FAO, and ameliorates steatosis without inflammatory or fibrotic liver injuries. More generally, our synthetic lipophagy adaptor protein, comprised of a LD-localizing sequence fused to a LIR, enables the selective analysis of LD autophagy, in contrast to bulk autophagy.

NAFLD begins with excessive lipid accumulation in the liver and, in some cases, progresses to NASH and liver cirrhosis ^13^. Hepatosteatosis is mainly caused by overnutrition, obesity, and insulin resistance. Other mechanisms subsequently contribute to liver injury, including oxidative stress, ER stress, mitochondrial dysfunction, as well as immunological alterations. The complex pathogenesis of NAFLD/NASH underscores the potential benefit of intervening in the initial steps of the disease, i.e., lipid accumulation. Lipid accumulation reflects a mis-balance between lipogenesis and lipolysis. Inhibiting lipogenesis can reduce steatosis without increasing inflammatory and profibrotic response, as seen for example in mice lacking hepatic Diacylglycerol Acyltransferase 2 (DGAT2)^50,51.^ Conversely, cytoplasmic lipolysis also decreases steatosis; however, in this case the ensuing production of cytoplasmic nonesterified fatty acids induces hepatic inflammation and systemic insulin resistance ^11,12.^ In contrast, the present study demonstrates that the lypolysis of LDs in autophagosomes and lysosomes via lipophagy safely disposes the lipid to the outside the liver and protects from lipotoxicity.

Previous efforts to promote liver lipophagy involved manipulating bulk autophagy. However, the LD autophagy was only a small fraction of total autophagy, even when maximally stimulated by starvation (Supplementary Fig. 3c). Our study is the first to examine the effect specifically of lipophagy on NAFLD/NASH pathology. To induce selective autophagy, we generated an adaptor protein composed of an LD-targeting sequence and an autophagosome recruiting domain. Similar adaptor approaches to achieve selective autophagy were previously reported using p62 ^52^, AMBRA1 ^26^ or small compounds ^53^ as LIR. We employed instead OPTN, which we found to have stronger affinity to autophagosome, and additionally introduced mutations to enhance the affinity against LC3. In addition, OPTN was trimmed to the minimal required sequence, to avoid unexpected effects mediated by other protein-protein interactions, as commonly seen with LC3 receptors ^27^. Thus, our newly engineered LIR is short, has strong capacity for autophagosome recruitment, and carries minimal risk of other effects. Our optimized approach can be extended to other selective autophagies with proper targeting domains, including mitophagy and pexophagy as shown here as well.

The canonical fate of autophagy is the degradation of cargo by lysosomal enzymes. Alternatively, autophagy can also have a secretory fate through exocytosis of autophagosomes or autolysosomes ^54^. The mechanisms underlying this fate decision remain largely unknown. Recent work showed that autophagosome-derived vesicles mainly contained RNA binding proteins, which controlled their secretion by interacting with LC3 and others ^55^. In the case of lipophagy, we found lipid content to be altered significantly in cultured medium, but not in isolated EVs, which can be derived from late-endosome and autophagosome, suggesting an alternative pathway of lipid exocytosis ^56^. The lipophagy-mediated improvement of steatosis was abrogated by inhibition of the fusion of autophagosome and lysosome, and the cell surface expression of the lysosomal protein LAMP1 was upregulated by lipophagy, a common marker of lysosomal exocytosis ^57^. These results collectively suggest that lysosomal exocytosis, rather than secretory autophagy, was responsible for lipophagy-mediated lipid secretion.

Lysosome secretion involves the migration of lysosomes from the perinuclear region to the cell surface and fusion with the cell membrane. It plays an important role in bone resorption, antigen presentation, and cell membrane repair ^58^. Lysosomal fusion with the cell membrane is mediated by Ca2+ signaling. Lysosomes store Ca2+ to a concentration similar to that in endoplasmic reticulum (ER) ^59^. Lysosomal Ca2+ is released through the TRPML1 Ca2+ channel, and the secreted Ca2+ is sensed by cell membrane-associated SYT7, which activates the SNARE complex to promote lipid bilayer fusion of lysosomes and plasma membrane ^60^. Starvation induces TRPML1-mediated Ca2+ efflux, possibly through mTOR inhibition ^42^, and a recent study reported that starvation-mediated autophagy of LDs exported fatty acid through lysosomal exocytosis, which required lipolysis of the TG cargo to activate the exocytosis ^61^. However, in our study, lipophagy was not accompanied by mTOR inhibition (Supplementary Fig. 6c); our Ca2+ assay suggested that fatty acids did not change the Ca2+ dynamics (Supplementary Fig. 6d); the lipid cargo was exported without lipase-mediated catabolization to nonesterified fatty acid (Fig. 3d); and lysosomal exocytosis was not activated by selective autophagy of peroxisome, despite their high content of fatty acids (Supplementary Fig. 6b). Thus, triglycerides, rather than fatty acids, might be involved in lysosomal deacidification and subsequent TRPML1-mediated Ca2+ efflux and shifting lysosome to exocytosis. One possibility is that loading too much cargo disturbs the trafficking of proton pump or ion channels, but the detailed mechanism of deacidification requires further investigation.

To identify potential inducers of lipophagy to treat NAFLD/NASH, we performed a small molecule library screen, using a newly developed lipophagy reporter system. ∼3500 clinically approved or functionally validated compounds were screened and several digitalis derivatives and PI3K-mTOR pathway inhibitors were identified as lipophagy activators. Among them, digoxin and a PI3K inhibitor, alpelisib were confirmed and showed significant therapeutic effects *in vivo* in a mouse model of NASH. Previous studies have reported the therapeutic potential of digoxin in NAFLD/NASH ^62,63.^ One study reported that digoxin binds pyruvate kinase M2 (PKM2) and inhibits chromatin remodeling, leading to downregulation of inflammatory HIF-1a transactivation ^62^. The other study identified cardiac glycosides as TFEB activators, via the known function of digoxin as an inhibitor of ATP-dependent Na+-K+ transporter, consequently activating Ca2+ influx through the sodium-calcium exchanger (NCX) or ER Ca2+ release by IP3R, in turn driving TFEB dephosphorylation and activation through calcineurin and an unknown phosphatase ^63,64^. In our study, unbiased screening identified digoxin as a lipophagy inducer, and we demonstrated *in vivo* that the protective effect of digoxin on the liver was entirely dependent on autophagy. However, digoxin treatment also improved the serum lipid profile, indicating that digoxin likely has additional liver effects on lipid metabolism. Digoxin has long been used clinically to modulate cardiac contractility and the conduction system. We took advantage of the prevalent use of digoxin to carry out a nested case control study, in which we demonstrated significantly lower prevalence of fatty liver in digoxin-treated patients. We conclude that digoxin provides a potential novel approach to protect against steatosis, although further clinical research is required.

In summary, our study demonstrates that a selective autophagy-inducing adaptor protein composed of a specific perilipin 1-derived LD-targeting domain fused to optimized LIR robustly activates lipophagy, safely disposes of aberrantly accumulated lipid through lysosomal exocytosis, and protects the liver from NAFLD/NASH in a mouse model. The system was also leveraged to establish a high-throughput screening system to identify lipophagy regulators and identify two FDA-approved drugs as lipophagy activators. The work provides multi-faceted proof-of-concept for the potential clinical application of lipophagy-based therapies.

## Acknowledgements

We thank the University of Pennsylvania Electron Microscopy Resource Laboratory for the support of electron microscopy study. We thank the University of Tokyo Drug Discovery Initiative for providing the compound library, which was supported by the Platform Project for Supporting Drug Discovery and Life Science Research from AMED (JP20am0101086, support number 2335). A.H. was supported by KAKENHI (19H03658), ONO Medical Research Foundation and Kato Memorial Bioscience Foundation.

## Author contributions

Y.M., A.H., Z.A., and S.M. designed the study. Y.M. executed most experiments with help from A.H., Y.H., Y.K., and S.T. Compound screening was performed by A.H. and Y.M. A case control study was done by M.H.and M.F. Y.K. analyzed tissue RNA-Seq. A.T. conducted Ca^2+^ imaging study. Y.M., A.H. and Z.A. wrote the manuscript. All authors discussed the results and commented on the manuscript.

## Materials availability

All unique/stable reagents generated in this study are available from the Lead Contact with a completed Materials Transfer Agreement

## Data availability

RNA-seq data have been deposited at GEO and are publicly available as of the date of publication; GSE185911 (https://www.ncbi.nlm.nih.gov/geo/query/acc.cgi?acc=GSE185911), token; itadoqsgxbwnpwd

## Methods

### Cells

HepG2 cells, 3T3L1 cells and Lenti-X 293T cells were cultured at 37 °C with 5% CO_2_ in Dulbecco’s modified Eagle’s medium (DMEM, Invitrogen) containing 10% fetal bovine serum (HyClone) and penicillin/streptomycin (100 U/ml, Invitrogen). To establish an NAFLD model in HepG2 cells, 0.1mM oleic acid (Sigma-Aldrich), or 0.5mM FFAs (oleic acid/ palmitic acid (Sigma-Aldrich), 2:1) for 24hr^65^. In some experiments, HepG2 cells were cultured with 25µM chloroquine (Sigma-Aldrich) for 24hr or 5nM Bafiromycin A1(Sigma-Aldrich) for 2hr^66^. HepG2 cells and 3T3L1 cells were purchased from ATCC, and Lenti-X 293T cells were from Clontech. No commonly misidentified cell line was used in this study. All the cell lines were routinely tested negative for mycoplasma contamination.

### Mice

All mouse experiments were approved by the Animal Care and Use Committee of the Kyoto Prefectural University of Medicine (M2020-59). C57BL/6J mice were purchased from CLEA Japan, Inc. B6.129S-Atg5<tm1Myok>(C57/BL6J background) were provided by the RIKEN BRC through the National Bio-Resource Project of the MEXT/AMED, Japan^67^. Chronic liver injury was induced by feeding a methionine/choline-deficient (MCD) diet (Research Diets Inc., A06071302) to 8-weeks-old male C57BL/6J or B6.129S-Atg5<tm1Myok> mice for 6 weeks. The MCD mice were grouped so that the average body weights of mice in each group were similar. The grouped mice were intravenously injected with a 2E11 genome copies (GC) /mouse of lipophagy induced (LDTD-LIR) or control (LDTD-dLIR) AAVs once, or orally administered digoxin (DG) (0.8mg/kg/day, Toronto Research Chemicals Inc, D446575), alpelisib (ALP) (50mg/kg/day, MedChem Express) or its solvent 0.5w/v% Methyl Cellulose solution (WAKO) once daily for two weeks^63,68.^ In some experiments, 65mg/kg chloroquine (Sigma-Aldrich) was administered by daily intraperitoneal injections. Mice were maintained in a specific pathogen-free animal facility on a 13:11 h light–dark cycle at an ambient temperature of 21 °C. They were given free access to water and food. Age- and sex-matched mice were used for all animal experiments.

### Human Study

We investigated a nested case-control study in a cohort study, which was performed in Kyoto Prefectural University of medicine. This cohort study was approved by Clinical Research Review Committee in Kyoto Prefectural University of medicine (CRB5200001, ERB-C-2104). We applied Opt-out method to obtain consent on this study. In this cohort, we extracted subjects who received digoxin and abdominal ultrasonography, as case subjects. Control subjects are matted with age, sex, BMI, diagnosis for atrial fibrillation. Fatty liver was diagnosed by the findings of abdominal ultrasonography performed by a trained technician. Among the four known diagnostic criteria; hepatorenal echo contrast, liver brightness, deep attenuation, and vascular blurring, hepatorenal echo contrast and liver brightness are required for fatty liver. The ultrasonographic definition is the same as those for NAFLD^49^.

### Palmitate Solution

Stock solutions were prepared as follows: palmitic acid (Sigma-Aldrich) was dissolved in 75% ethanol at 70 °C at a final concentration of 300 mM. Aliquots of stock solutions were complexed with fatty-acid–free BSA (10% solution in 150 mM NaCl; Sigma-Aldrich) by stirring for 1 h at 37 °C. The final molar ratio of fatty acid:BSA was 5:1. The final ethanol concentration of stock solution was 1.5% (vol:vol). All control conditions included a solution of vehicle (ethanol:H2O) mixed with fatty-acid–free BSA in NaCl solution at the same concentration as the palmitate solution^69^.

### Virus production

To produce viruses, 6-well plates of 70% confluent Lenti-X 293T(Clontech) cells were transfected with 1.5 μg of transfer vector, 0.5 μg pMD2.G and 1.0 μg of psPAX2 for lentivirus or 1.0 μg of gag/pol for retrovirus using Fugene HD(Promega) according to the manufacturer’s instructions. Supernatant was collected after 48 h and frozen at –80 °C.

### Electron microscopy

For EM studies, cultured 3T3L1 adipocytes and HepG2 hepatocytes were trypsinized, washed and fixed with 2.5% (vol/vol) glutaraldehyde, 2.0% (vol/vol) paraformaldehyde in 0.1 M sodium cacodylate buffer, pH 7.4, overnight at 4 °C. EM studies were performed on a JEM-1010 microscope at the University of Pennsylvania Electron Microscopy Resource Laboratory.

### Oil red O staining

The Oil Red O staining was performed by fixing HepG2 cells in 4% paraformaldehyde and then staining with Oil Red O for 15 min. The samples were washed with 60% isopropanol for a few seconds, followed by three PBS washes. Analysis of stain-positive regions was performed using image J (National Institutes of Health, Bethesda, MD).

### Histopathological analysis

Mice were killed and liver, kidney, skeletal muscle and WAT were fixed with 4% PFA overnight. Then fixed tissues were embedded in OCT or paraffin. The paraffin sections were used for hematoxylin and eosin (HE) staining and picrosirius red staining. The OCT sections were used for oil red O (Sigma-Aldrich, O0625) staining and immunohistochemistry (IHC) staining of αSMA. Analysis of stain-positive regions and adipocyte size was performed using ImageJ (National Institutes of Health, Bethesda, MD).

### RNA sequence

Total RNA was isolated from the livers using TRIzol (Life Technologies) and Direct-zol RNA MiniPrep (Zymo Research Corporation) according to the manufacturer’s protocol. The Library preparation was performed using a TruSeq stranded mRNA sample prep kit (Illumina) according to the manufacturer’s instructions. Sequencing was performed on an Illumina NOVASeq 6000 platform in a 100 bp paired-end mode.

### Real-time PCR

RNA was isolated from livers using TRIzol (Life Technologies) and Direct-zol RNA MiniPrep (Zymo Research Corporation) according to the manufacturer’s instructions. Using the PrimeScript RT Master Mix (Takara Bio), we reverse transcribed total RNA. The cDNA was amplified by primers in a 10µl reaction using KAPA SYBR FAST (Kapa Biosystems). We calculated mRNA using a ΔΔCT relative to the average of the housekeeping genes GAPDH expression. All primers and gene information were provided in Table S1.

### Immunoblot

Total protein concentration of cell or liver lysate was determined using the Lowry assay (Bio-Rad Laboratories, 5000112JA). Equal amounts of protein were loaded onto Tris-glycine sodium dodecyl sulfate-polyacrylamide gels, separated by electrophoresis, and then the proteins were transferred onto polyvinylidene difluoride membranes (Millipore, IPVH00010). Membranes were then probed with anti-α-SMA (Abcam, ab5694, 1:100 for IHC, 1:1000 for WB), anti-ATG5 (Santa Cruz Biotechnology, sc133158, 1:500 for WB), anti-S6 Ribosomal Protein (CST, 2217, 1:1000 for WB), anti-Phospho-S6 Ribosomal Protein (Ser235/236) (CST, 2211, 1:1000 for WB), and anti-β-actin antibodies followed by incubation with a HRP-conjugated secondary antibody. Immunolabeled bands were detected by chemiluminescence using the Clarity Western ECL substrate (Bio-Rad Laboratories,1705060) and the Clarity MAX Western ECL substrate (Bio-Rad Laboratories,1705062). Densitometric analysis of band intensity was performed using ImageJ (National Institutes of Health, Bethesda, MD).

### Biochemical assays of tissues

Lipids of tissue, serum exosome, cell lysate and supernatant were extracted by the Folch method^70^. Triglyceride (TG) and non-esterified fatty acid (NEFA) concentrations were measured using test kits (LabAssay(tm) Triglyceride Kit, LabAssay(tm) NEFA Kit: Wako), according to the manufacturer’s instructions.

### Serum Lipid Profiling

Cholesterol and triglycerides in lipoproteins were analyzed by HPLC at Skylight Biotech (Akita, Japan)^71^.

### Fatty acid oxidation (FAO) activity assay

FAO enzyme activities were measured using FAO Assay Kit (Biomedical Research Service Center, E-141), according to the manufacturer’s instructions^72^. All samples were harvested using 1× Cell Lysis Solution. Protein concentration of the samples was assessed with a Lowry assay and normalized to 1 mg/ml. Add 20 µl of each sample to a plain 96-well plate placed on ice in duplicate. Then, swiftly add 50 µl control solution to one set of wells and 50 µl reaction solution to the other set of wells. Mix contents by gentle agitation for 10 sec. Cover plate and keep in a non-CO2 incubator at 37°C for 30 min. The plate was read at optical density of 492 nm (OD 492) with a microplate reader (BIO-RAD, iMark). Subtract control well reading from reaction well reading for each sample. FAO activity in IU/l unit was determined by multiplying OD by 12.96.

### Extracellular vesicles isolation

Exosome isolation from serum sample was performed using MagCapture Exosome Isolation Kit PS (Wako, 293-77601), according to the manufacturer’s instructions^73^. Magnetic beads coated with Tim4 protein (which binds to Phosphatidylserine on the membrane surface of extracellular vesicles) enable purification of highly purified EVs.

### BODIPY staining

For BODIPY 494/503 (4,4-Difluoro-1,3,5,7,8-Pentamethyl-4-Bora-3a,4a-Diaza-s-Indacene, Thermo Fisher Scientific, D3922) staining, cells were washed and stained with 2µm BODIPY staining solution for 15 min at 37°C. Cells were then washed, harvested by trypsinization and resuspended with FACS buffer (PBS containing 2% FBS and 20 mM HEPES). Stained cells were analyzed on Attune NxT Flow Cytometer (Thermo Fisher Scientific) and analysis was done with FlowJo (Treestar)^74^.

### LysoSensor

For LysoSensor Green DND-189 (Thermo Fisher Scientific, L7535) staining, cells were washed and stained with 50nM LysoSensor for 30min at 37°C. Cells were then washed, harvested by trypsinization and resuspended with FACS buffer (PBS containing 2% FBS and 20 mM HEPES). Stained cells were analyzed on Attune NxT Flow Cytometer (Thermo Fisher Scientific) and analyzed with FlowJo (Treestar)^75^.

### Lysosomal acid lipase assay

This assay was performed using LysoLive Lysosomal Acid Lipase Assay Kit (Abcam, ab253380). Briefly, lipophagy-induced HepG2 cells were incubated LipaGreen for 8 hr at 37°C. Cells were then washed, harvested by trypsinization and resuspended with FACS buffer (PBS containing 2% FBS and 20 mM HEPES). Stained cells were analyzed on Attune NxT Flow Cytometer (Thermo Fisher Scientific) and analyzed with FlowJo (Treestar).

### Lysosomal exocytosis analysis

To assess lysosomal exocytosis, we used LAMP1 expression on the membrane surface^76^. Cells were harvested by trypsinization, washed and incubated with anti-LAMP1 antibodies (Biolegend, 328606, 1:50) for 20min on ice. Cells were then washed and resuspended with FACS buffer. Stained cells were analyzed on Attune NxT Flow Cytometer (Thermo Fisher Scientific) and analysis was done with FlowJo (Treestar).

### Fura-2 Ca^2+^ imaging

293T cells expressing TRPML1 on the membrane surface were prepared, and were plated onto glass coverslips. Cells were loaded with 5µM Fura-2 AM (DOJINDO, 343-05401) in the culture medium at 37°C for 60min. The extracellular bath solution (Tyrode’s solution) contained 153mM NaCl, 5mM KCl, 2mM CaCl2, 1mM MgCl2, 20mM HEPES and 10mM glucose (pH 7.4). The Low pH Tyrode solution contained 150mM Na-gluconate, 5mM KCl, 2mM CaCl2, 1mM MgCl2, 10mM glucose, 10mM HEPES and 10mM MES (pH 4.6). Oleic acid (Sigma-Aldrich) and fatty acid mixture (Sigma-Aldrich, L0288) were used as fatty acids. ML-SA1 (10µM, TRPML agonist, Wako, 131-18531) were used as positive controls to induce Ca^2+^ release from lysosome and acidic stores. All bath solutions were applied via a perfusion system to achieve a complete solution exchange within a few seconds. Cells were recorded on the stage of an inverted microscope (IX-73, Olympus, Tokyo, Japan; 20 × 0.7 NA UCPlanFLN20 x PH) with continuous perfusion (RC-27L, Warner Instruments, Hamden, CT, USA). Cytosolic free Ca^2+^ concentrations were measured by dual-wavelength Fura-2 microfluorometry with excitation at 340/380 nm and emission at 510 nm. The ratio image was calculated and acquired using an sCMOS camera (ORCA-Flash4.0, Hamamatsu Photonics, Tokyo, Japan) and the HCImage software (Hamamatsu Photonics) ^77^.

### Intraperitoneal glucose tolerance test (IPGTT) and intraperitoneal insulin tolerance test (IPITT)

IPGTT and ITT were performed on 16 weeks old mice two weeks after injection of AAV. After fasting, the baseline blood glucose level was measured by tail vein puncture. For IPGTT, mice were fasted for 15 h and a solution of 20% glucose (2g/kg body weight) was administered intraperitoneally. After glucose administration, blood samples were collected from the tail vein at 15, 30, 60, and 120 min. For IPITT, mice were fasted for 5 h. Insulin-R(Eli-Lilly) was intraperitoneally injected (1U/kg body weight) and blood samples from the tail vein were collected at 15, 30, 60, and 120 min after insulin injection. Glucose levels were evaluated with Glutentmint (Sanwa Kagaku Kenkyusho)^69^.

### Fasting Plasma Insulin and Homeostatic Model Assessment of Insulin Resistance (HOMA-IR)

The blood samples collected after 15 h of fasting were used for the quantification of plasma insulin level with an enzyme-linked immunosorbent assay (ELISA), according to the manufacturer’s recommendations (MIoBS, M1104). HOMA-IR was estimated from fasting glucose and insulin as follows:

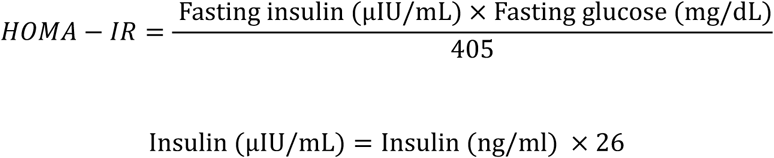

### Compound library screening

HepG2 cells expressing GFP-mCherry-LDTS and nuclear TagBFP were generated as lipophagy reporter cells. Clinically-approved or validated ∼3500 compound library was provided as Plate on Demand (POD) by Drug Discovery Initiative in The University of Tokyo. Total of 5000 reporter cells were seeded by Multidrop 384 dispenser (ThermoFisher) in POD 384 well plate. After 24 hr-culture, cells were analyzed with IN Cell Analyzer 2200. IN Cell Analyzer workstation was used to count TagBFP positive dots as cell number and measure total fluorescence of GFP and mCherry in mCherry-positive regions. Lipophagy score was calculated by (mCherry total fluorescence - GFP total fluorescence) in mCherry positive area and normalized with corresponding cell number. A DMSO-treated plate was used to correct position artifacts. For each plate, the raw value of lipophagy score was converted according to the value of DMSO-treated plat for all wells. Robust Z-scores were calculated using the following equation.

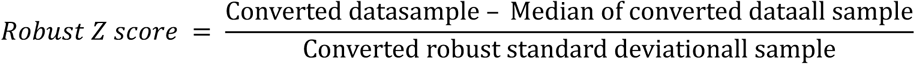

### Statistical analyses

All data were expressed as mean ± s.d. No statistical methods were used to predetermine sample size. Sample size was based on experimental feasibility and sample availability. Samples were processed in random order. Comparisons between the two groups were analysed using the two-sided unpaired t-test. One-way ANOVA followed by Tukey’s post hoc test was used for multiple group comparisons.; P < 0.05 was considered statistically significant.

## References

1. Rinella, M. & Charlton, M. The globalization of nonalcoholic fatty liver disease: Prevalence and impact on world health. Hepatology 64, 19–22 (2016).

2. Younossi, Z.M., et al. Global epidemiology of nonalcoholic fatty liver disease-Meta-analytic assessment of prevalence, incidence, and outcomes. Hepatology 64, 73–84 (2016).

3. Tilg, H. & Moschen, A.R. Evolution of inflammation in nonalcoholic fatty liver disease: the multiple parallel hits hypothesis. Hepatology 52, 1836–1846 (2010).

4. Angulo, P., et al. Liver Fibrosis, but No Other Histologic Features, Is Associated With Long-term Outcomes of Patients With Nonalcoholic Fatty Liver Disease. Gastroenterology 149, 389–397 e310 (2015).

5. Krenkel, O., et al. Therapeutic inhibition of inflammatory monocyte recruitment reduces steatohepatitis and liver fibrosis. Hepatology 67, 1270–1283 (2018).

6. Marcher, A.B., et al. Transcriptional regulation of Hepatic Stellate Cell activation in NASH. Sci Rep 9, 2324 (2019).

7. Pydyn, N., Miekus, K., Jura, J. & Kotlinowski, J. New therapeutic strategies in nonalcoholic fatty liver disease: a focus on promising drugs for nonalcoholic steatohepatitis. Pharmacol Rep 72, 1–12 (2020).

8. Ratziu, V., Goodman, Z. & Sanyal, A. Current efforts and trends in the treatment of NASH. J Hepatol 62, S65–75 (2015).

9. Sanyal, A.J., et al. Challenges and opportunities in drug and biomarker development for nonalcoholic steatohepatitis: findings and recommendations from an American Association for the Study of Liver Diseases-U.S. Food and Drug Administration Joint Workshop. Hepatology 61, 1392–1405 (2015).

10. Sztalryd, C. & Brasaemle, D.L. The perilipin family of lipid droplet proteins: Gatekeepers of intracellular lipolysis. Biochim Biophys Acta Mol Cell Biol Lipids 1862, 1221–1232 (2017).

11. Wang, C., et al. Perilipin 5 improves hepatic lipotoxicity by inhibiting lipolysis. Hepatology 61, 870–882 (2015).

12. Trevino, M.B., et al. Liver Perilipin 5 Expression Worsens Hepatosteatosis But Not Insulin Resistance in High Fat-Fed Mice. Mol Endocrinol 29, 1414–1425 (2015).

13. Mashek, D.G. Hepatic lipid droplets: A balancing act between energy storage and metabolic dysfunction in NAFLD. Mol Metab, 101115 (2020).

14. Czaja, M.J. & Cuervo, A.M. Lipases in lysosomes, what for? Autophagy 5, 866–867 (2009).

15. Burke, J.A. & Schubert, W.K. Deficient activity of hepatic acid lipase in cholesterol ester storage disease. Science 176, 309–310 (1972).

16. Patrick, A.D. & Lake, B.D. Deficiency of an acid lipase in Wolman’s disease. Nature 222, 1067–1068 (1969).

17. Singh, R., et al. Autophagy regulates lipid metabolism. Nature 458, 1131–1135 (2009).

18. Schulze, R.J., Sathyanarayan, A. & Mashek, D.G. Breaking fat: The regulation and mechanisms of lipophagy. Biochim Biophys Acta Mol Cell Biol Lipids 1862, 1178–1187 (2017).

19. Singh, R., et al. Autophagy regulates adipose mass and differentiation in mice. J Clin Invest 119, 3329–3339 (2009).

20. Zhang, Y., et al. Adipose-specific deletion of autophagy-related gene 7 (atg7) in mice reveals a role in adipogenesis. Proc Natl Acad Sci U S A 106, 19860–19865 (2009).

21. Ma, D., et al. Autophagy deficiency by hepatic FIP200 deletion uncouples steatosis from liver injury in NAFLD. Mol Endocrinol 27, 1643–1654 (2013).

22. Ramos, V.M., Kowaltowski, A.J. & Kakimoto, P.A. Autophagy in Hepatic Steatosis: A Structured Review. Front Cell Dev Biol 9, 657389 (2021).

23. Settembre, C., et al. TFEB controls cellular lipid metabolism through a starvation-induced autoregulatory loop. Nat Cell Biol 15, 647–658 (2013).

24. Tanaka, S., et al. Rubicon inhibits autophagy and accelerates hepatocyte apoptosis and lipid accumulation in nonalcoholic fatty liver disease in mice. Hepatology 64, 1994–2014 (2016).

25. Katayama, H., Yamamoto, A., Mizushima, N., Yoshimori, T. & Miyawaki, A. GFP-like proteins stably accumulate in lysosomes. Cell Struct Funct 33, 1–12 (2008).

26. Strappazzon, F., et al. AMBRA1 is able to induce mitophagy via LC3 binding, regardless of PARKIN and p62/SQSTM1. Cell Death Differ 22, 517 (2015).

27. Ryan, T.A. & Tumbarello, D.A. Optineurin: A Coordinator of Membrane-Associated Cargo Trafficking and Autophagy. Front Immunol 9, 1024 (2018).

28. Heo, J.M., Ordureau, A., Paulo, J.A., Rinehart, J. & Harper, J.W. The PINK1-PARKIN Mitochondrial Ubiquitylation Pathway Drives a Program of OPTN/NDP52 Recruitment and TBK1 Activation to Promote Mitophagy. Mol Cell 60, 7–20 (2015).

29. Rogov, V.V., et al. Structural basis for phosphorylation-triggered autophagic clearance of Salmonella. Biochem J 454, 459–466 (2013).

30. Tanenbaum, M.E., Gilbert, L.A., Qi, L.S., Weissman, J.S. & Vale, R.D. A protein-tagging system for signal amplification in gene expression and fluorescence imaging. Cell 159, 635–646 (2014).

31. Tanaka, T., et al. Direct targeting of proteins to lipid droplets demonstrated by time-lapse live cell imaging. J Biosci Bioeng 116, 620–623 (2013).

32. Murugesan, S., Goldberg, E.B., Dou, E. & Brown, W.J. Identification of diverse lipid droplet targeting motifs in the PNPLA family of triglyceride lipases. PLoS One 8, e64950 (2013).

33. Garcia, A., Sekowski, A., Subramanian, V. & Brasaemle, D.L. The central domain is required to target and anchor perilipin A to lipid droplets. J Biol Chem 278, 625–635 (2003).

34. Chiba, T., Suzuki, S., Sato, Y., Itoh, T. & Umegaki, K. Evaluation of Methionine Content in a High-Fat and Choline-Deficient Diet on Body Weight Gain and the Development of Non-Alcoholic Steatohepatitis in Mice. PLoS One 11, e0164191 (2016).

35. Mauthe, M., et al. Chloroquine inhibits autophagic flux by decreasing autophagosome-lysosome fusion. Autophagy 14, 1435–1455 (2018).

36. Uchinomiya, S., et al. Fluorescence detection of metabolic activity of the fatty acid beta oxidation pathway in living cells. Chem Commun (Camb) 56, 3023–3026 (2020).

37. Huynh, C., Roth, D., Ward, D.M., Kaplan, J. & Andrews, N.W. Defective lysosomal exocytosis and plasma membrane repair in Chediak-Higashi/beige cells. Proc Natl Acad Sci U S A 101, 16795–16800 (2004).

38. Fransen, M., Wylin, T., Brees, C., Mannaerts, G.P. & Van Veldhoven, P.P. Human pex19p binds peroxisomal integral membrane proteins at regions distinct from their sorting sequences. Mol Cell Biol 21, 4413–4424 (2001).

39. Gould, S.J., Keller, G.A., Hosken, N., Wilkinson, J. & Subramani, S. A conserved tripeptide sorts proteins to peroxisomes. J Cell Biol 108, 1657–1664 (1989).

40. Samie, M., et al. A TRP channel in the lysosome regulates large particle phagocytosis via focal exocytosis. Dev Cell 26, 511–524 (2013).

41. Flannery, A.R., Czibener, C. & Andrews, N.W. Palmitoylation-dependent association with CD63 targets the Ca2+ sensor synaptotagmin VII to lysosomes. J Cell Biol 191, 599–613 (2010).

42. Wang, W., et al. Up-regulation of lysosomal TRPML1 channels is essential for lysosomal adaptation to nutrient starvation. Proc Natl Acad Sci U S A 112, E1373–1381 (2015).

43. Shen, D., et al. Lipid storage disorders block lysosomal trafficking by inhibiting a TRP channel and lysosomal calcium release. Nat Commun 3, 731 (2012).

44. Zhou, J., et al. Activation of lysosomal function in the course of autophagy via mTORC1 suppression and autophagosome-lysosome fusion. Cell Res 23, 508–523 (2013).

45. Ameis, D., Merkel, M., Eckerskorn, C. & Greten, H. Purification, characterization and molecular cloning of human hepatic lysosomal acid lipase. Eur J Biochem 219, 905–914 (1994).

46. Albrecht, L.V., Tejeda-Munoz, N. & De Robertis, E.M. Protocol for Probing Regulated Lysosomal Activity and Function in Living Cells. STAR Protoc 1, 100132 (2020).

47. Miao, Y., Li, G., Zhang, X., Xu, H. & Abraham, S.N. A TRP Channel Senses Lysosome Neutralization by Pathogens to Trigger Their Expulsion. Cell 161, 1306–1319 (2015).

48. Stahl, E.P., et al. Nonalcoholic Fatty Liver Disease and the Heart: JACC State-of-the-Art Review. J Am Coll Cardiol 73, 948–963 (2019).

49. Hamaguchi, M., et al. The severity of ultrasonographic findings in nonalcoholic fatty liver disease reflects the metabolic syndrome and visceral fat accumulation. Am J Gastroenterol 102, 2708–2715 (2007).

50. Gluchowski, N.L., et al. Hepatocyte Deletion of Triglyceride-Synthesis Enzyme Acyl CoA: Diacylglycerol Acyltransferase 2 Reduces Steatosis Without Increasing Inflammation or Fibrosis in Mice. Hepatology 70, 1972–1985 (2019).

51. Loomba, R., et al. Novel antisense inhibition of diacylglycerol O-acyltransferase 2 for treatment of non-alcoholic fatty liver disease: a multicentre, double-blind, randomised, placebo-controlled phase 2 trial. Lancet Gastroenterol Hepatol 5, 829–838 (2020).

52. Tatsumi, T., et al. Forced lipophagy reveals that lipid droplets are required for early embryonic development in mouse. Development 145(2018).

53. Takahashi, D., et al. AUTACs: Cargo-Specific Degraders Using Selective Autophagy. Mol Cell 76, 797–810 e710 (2019).

54. Nieto-Torres, J.L., Leidal, A.M., Debnath, J. & Hansen, M. Beyond Autophagy: The Expanding Roles of ATG8 Proteins. Trends Biochem Sci (2021).

55. Leidal, A.M., et al. The LC3-conjugation machinery specifies the loading of RNA-binding proteins into extracellular vesicles. Nat Cell Biol 22, 187–199 (2020).

56. Xu, J., Camfield, R. & Gorski, S.M. The interplay between exosomes and autophagy-partners in crime. J Cell Sci 131(2018).

57. Andrews, N.W. Detection of Lysosomal Exocytosis by Surface Exposure of Lamp1 Luminal Epitopes. Methods Mol Biol 1594, 205–211 (2017).

58. Tancini, B., et al. Lysosomal Exocytosis: The Extracellular Role of an Intracellular Organelle. Membranes (Basel) 10(2020).

59. Lopez Sanjurjo, C.I., Tovey, S.C. & Taylor, C.W. Rapid recycling of Ca2+ between IP3-sensitive stores and lysosomes. PLoS One 9, e111275 (2014).

60. Czibener, C., et al. Ca2+ and synaptotagmin VII-dependent delivery of lysosomal membrane to nascent phagosomes. J Cell Biol 174, 997–1007 (2006).

61. Cui, W., et al. Lipophagy-derived fatty acids undergo extracellular efflux via lysosomal exocytosis. Autophagy, 1–16 (2020).

62. Ouyang, X., et al. Digoxin Suppresses Pyruvate Kinase M2-Promoted HIF-1alpha Transactivation in Steatohepatitis. Cell Metab 27, 339–350 e333 (2018).

63. Wang, C., et al. Small-molecule TFEB pathway agonists that ameliorate metabolic syndrome in mice and extend C. elegans lifespan. Nat Commun 8, 2270 (2017).

64. Tesselaar, M.H., et al. Digitalis-like Compounds Facilitate Non-Medullary Thyroid Cancer Redifferentiation through Intracellular Ca2+, FOS, and Autophagy-Dependent Pathways. Mol Cancer Ther 16, 169–181 (2017).

65. Dhami-Shah, H., et al. Picroside II attenuates fatty acid accumulation in HepG2 cells via modulation of fatty acid uptake and synthesis. Clin Mol Hepatol 24, 77–87 (2018).

66. Yan, Y., et al. Bafilomycin A1 induces caspase-independent cell death in hepatocellular carcinoma cells via targeting of autophagy and MAPK pathways. Sci. Rep. 6, 37052 (2016).

67. Hara, T., et al. Suppression of basal autophagy in neural cells causes neurodegenerative disease in mice. Nature 441, 885–889 (2006).

68. Gobin, B., et al. BYL719, a new alpha-specific PI3K inhibitor: single administration and in combination with conventional chemotherapy for the treatment of osteosarcoma. Int. J. Cancer 136, 784–796 (2015).

69. Hoshino, A., et al. Inhibition of p53 preserves Parkin-mediated mitophagy and pancreatic beta-cell function in diabetes. Proc. Natl. Acad. Sci. U. S. A. 111, 3116–3121 (2014).

70. Folch, J., Lees, M. & Stanley, G.H.S. A Simple Method for the Isolation and Purification of Total Lipides from Animal Tissues. J. Biol. Chem. 226, 497–509 (1957).

71. Usui, S., Hara, Y., Hosaki, S. & Okazaki, M. A new on-line dual enzymatic method for simultaneous quantification of cholesterol and triglycerides in lipoproteins by HPLC. J Lipid Res 43, 805–814 (2002).

72. Kwong, S.C., Jamil, A.H.A., Rhodes, A., Taib, N.A. & Chung, I. Metabolic role of fatty acid binding protein 7 in mediating triple-negative breast cancer cell death via PPAR-alpha signaling. J. Lipid Res. 60, 1807–1817 (2019).

73. Liu, Z., et al. Exosomes from adipose-derived mesenchymal stem cells prevent cardiomyocyte apoptosis induced by oxidative stress. Cell Death Discov 5, 79 (2019).

74. Qiu, B. & Simon, M.C. BODIPY 493/503 Staining of Neutral Lipid Droplets for Microscopy and Quantification by Flow Cytometry. Bio Protoc 6(2016).

75. Ghosh, S., et al. beta-Coronaviruses Use Lysosomes for Egress Instead of the Biosynthetic Secretory Pathway. Cell 183, 1520–1535 e1514 (2020).

76. Samie, M.A. & Xu, H. Lysosomal exocytosis and lipid storage disorders. J. Lipid Res. 55, 995–1009 (2014).

77. Shen, D., et al. Lipid storage disorders block lysosomal trafficking by inhibiting a TRP channel and lysosomal calcium release. Nature Communications 3(2012).

