## Supplementary data for "Liver lipophagy ameliorates nonalcoholic steatohepatitis through lysosomal lipid exocytosis"

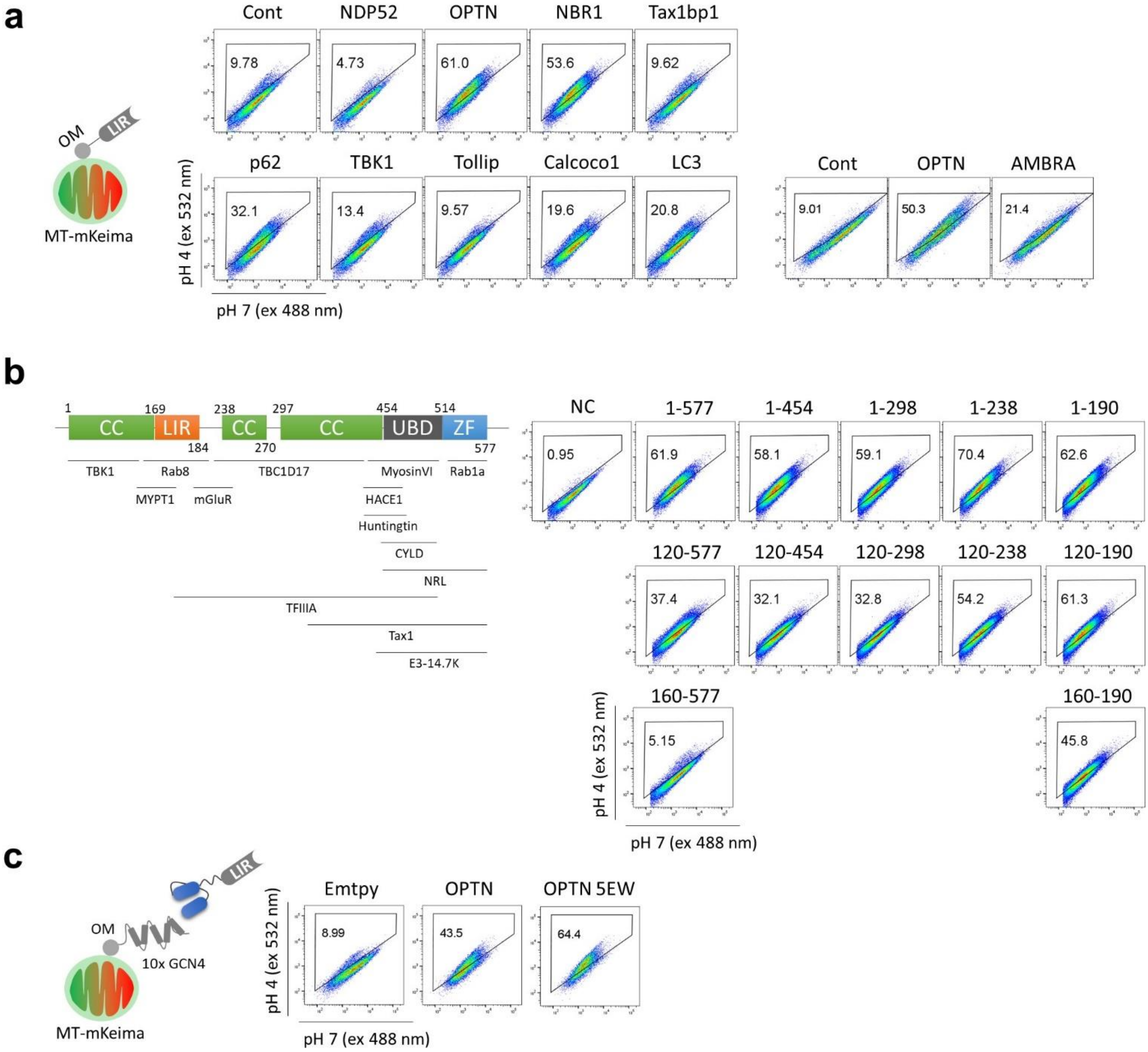


**Supplementary Figure 1. Engineering LIR domain.**

(**a**) Autophagosome recruiting capacity of LC3 receptors assessed by mitophagy induction. Mitophagy level was quantified by mt-mKeima. (**b**) Domains of optineurin (OPTN) and interacting regions with binding partners (left). Identification of essential sequence for autophagosome recruiting capability (right). (**c**) Autophagosome recruiting capacity of phosphomimetic and tryptophan mutation (5EW) in full-length of OPTN in the SunTag system.

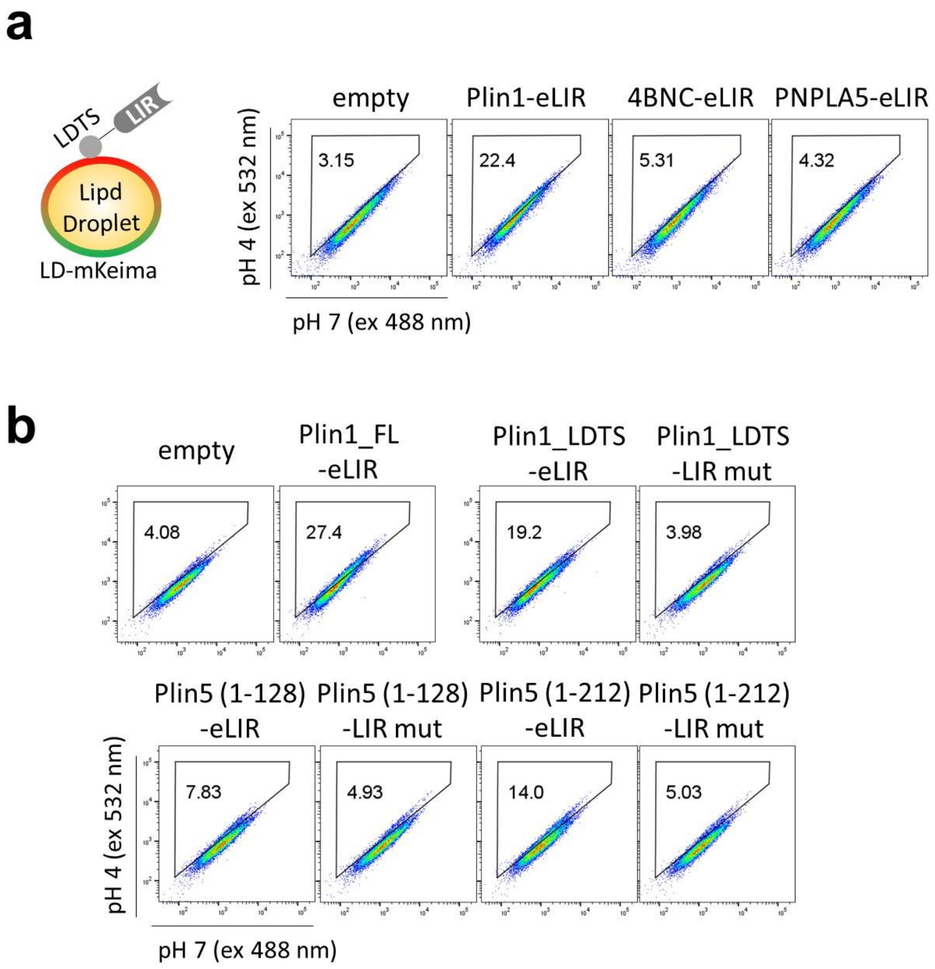


**Supplementary Figure 2. Optimization of lipid droplet targeting signal.**

(**a**) Lipid droplet (LD) targeting capability of LD-related protein assessed by lipophagy induction in the fusion of engineered LIR (eLIR). (**b**) Essential sequence if LD-targeting signal (LDTS) of perilipin 1 and perilipin 5.

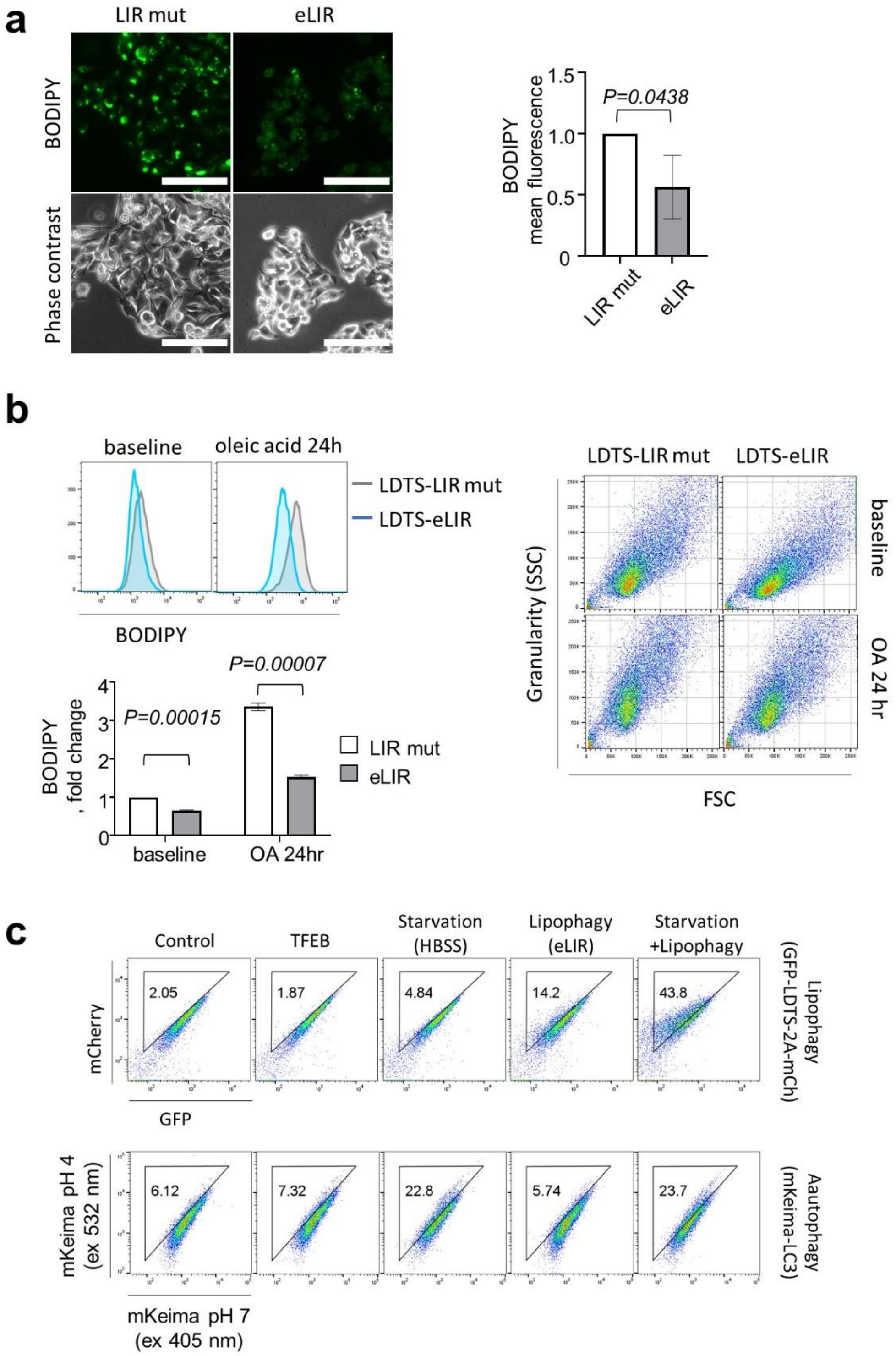


**Supplementary Figure 3. Massive induction of lipophagy with LDTS-eLIR.**

(**a**) Fluorescence images of HepG2 cells stained with BODIPY after 24-hour incubation with oleic acid. induction. Scale bars, 100 µm. (**b**) Quantification of BODIPY in HepG2 cells by flow cytometry (Left), and the intracellular granularity reflecting the content of lipid droplets (right). (**c**) The extent of lipophagy induction and concomitant autophagy activation with TFEB overexpression, starvation, and LDTS-eLIR-mediated lipophagy. All data are presented as mean±SD of n=3. *P* values calculated by two-sided unpaired *t*-test.

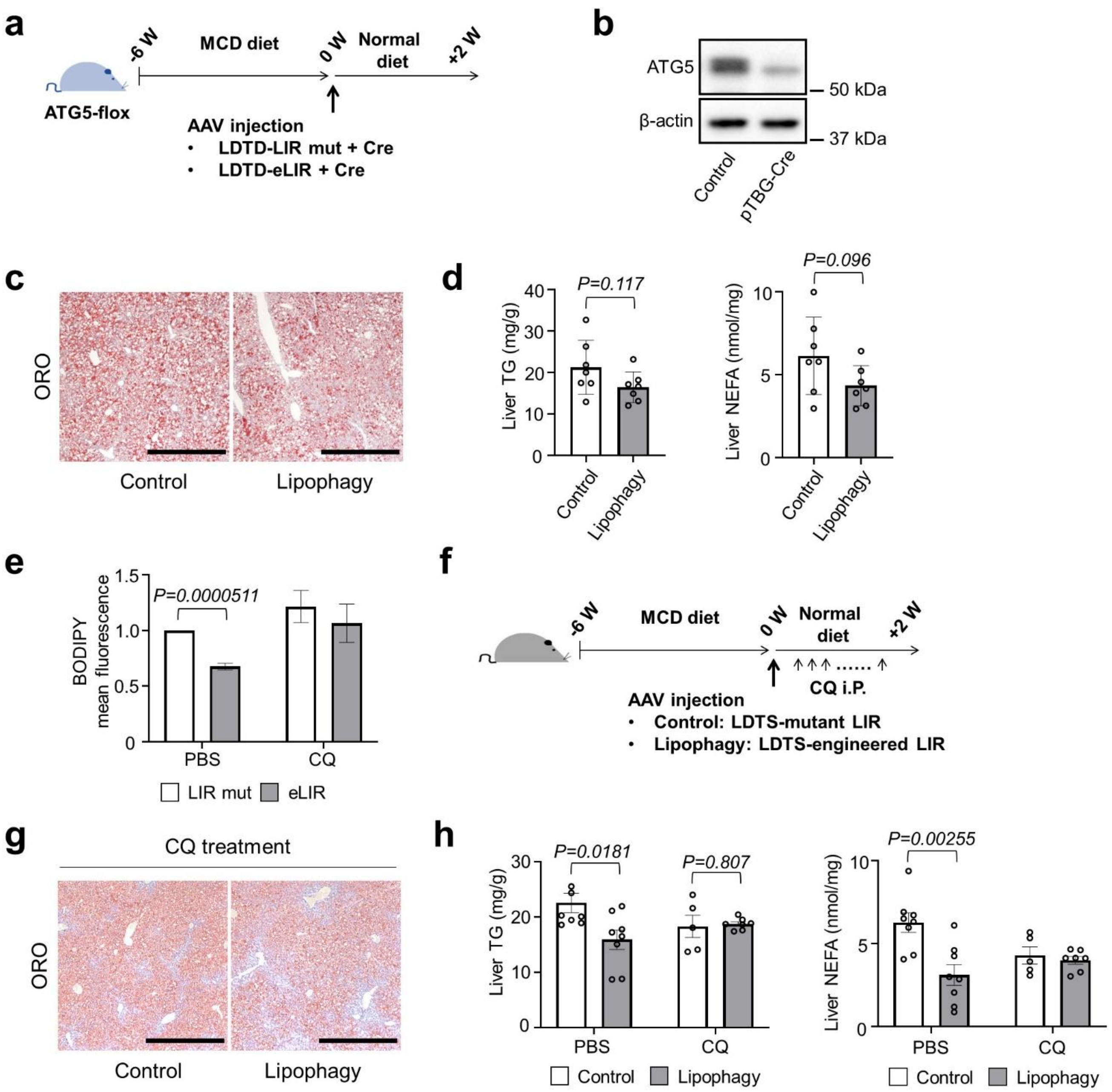


**Supplementary Figure 4. Liver-specific ATG5 deletion and chloroquine treatment cancelled the effects of lipophagy.**

(**a**) Study protocol of NASH model with MCD diet, liver-specific ATG5 knockout and lipophagy induction. (**b**) Immunoblots assessing the expression of ATG5 in liver tissue isolated from ATG5-flox mice injected with AAV8 pTBG-Cre. (**c**) Oil red O staining of liver sections. Scale bars, 1 mm. (**d**) Liver TG and NEFA content. (n=7 for each group). (**e**) Quantification of BODIPY by flow cytometry in HepG2 cells treated with chloroquine (CQ, 25μM) and oleic acid for 24 hours (n=3). (**f**) Study protocol of NASH model with MCD diet, CQ treatment and lipophagy induction. (**g**) ORO staining of liver sections. Scale bars, 1000μm. (**h**) Liver TG and NEFA content. (n=8 CQ- groups, n=5 CQ+/control, n=7 CQ+/lipophagy). All data are presented as mean±SD. *P* values calculated by two-sided unpaired *t*-test.

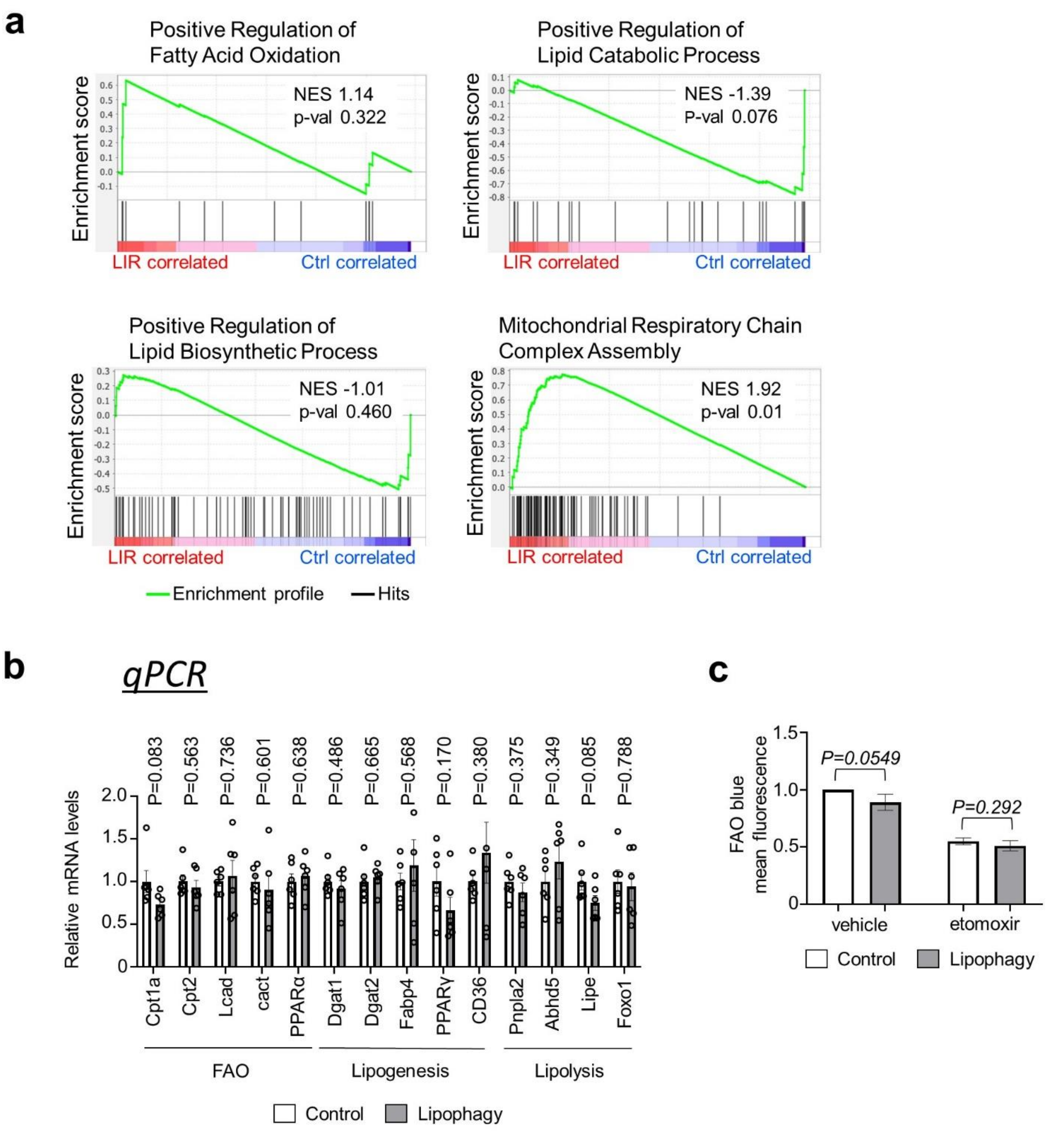


**Supplementary Figure 5. Lipid metabolism in lipophagy-induced livers and HepG2 cells.**

(**a**) Gene set enrichment analysis for lipid metabolism in the RNA sequence of liver tissues related with Figure 2i,j. (**b**) Real-time quantitative PCR assessing the expression of genes associated with lipid metabolism in liver tissues. Data are presented as mean±SD of n=6 per group. (**c**) FAO activity in the cultured HepG2 cells. FAO blue is quenched fluorescent probe and metabolically activated by the sequential enzyme reactions of FAO. Data are presented as mean±SD of n=3. *P* values calculated by two-sided unpaired *t*-test.

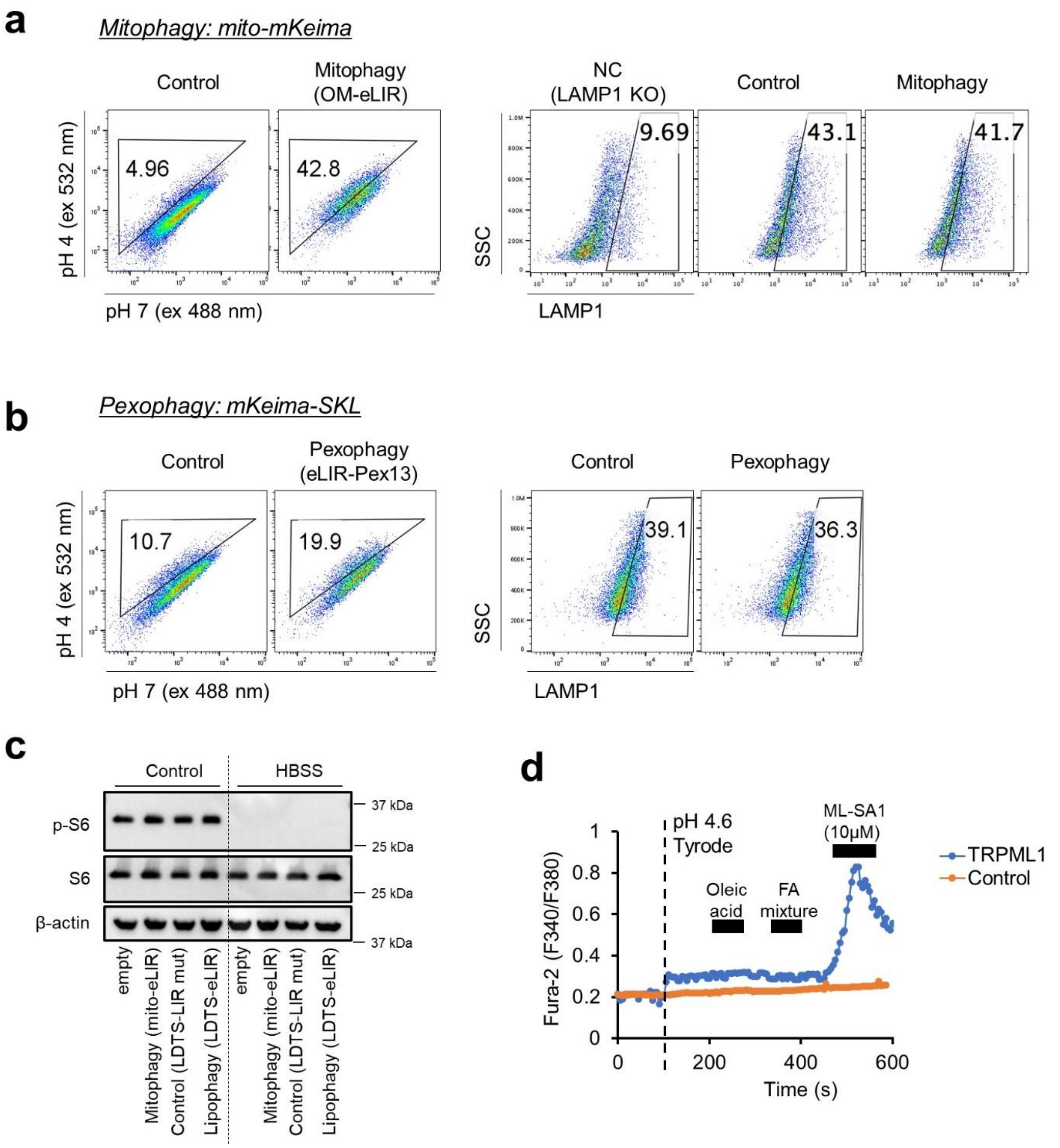


**Supplementary Figure 6. Lysosomal exocytosis is independent of mTOR1 signal and nonesterified fatty acid.** (**a**) Mitophagy induction by mitochondrial outermembrane-targeted eLIR was confirmed by flow cytometry for mt-mKeima in HepG2 cells (left). LAMP1 in the surface of HepG2 cells after induction of mitophagy (right). LAMP1 KO cells were used as a negative control. (**b**) Pexophagy induction by fusion protein of the localization motif of the peroxin Pex13 and eLIR was confirmed by flow cytometry for peroxisome-localized mKeima-SKL in HepG2 cells (left). LAMP1 in the surface of HepG2 cells after induction of pexophagy (right). (**c**) WB in lipophagy-induced cell under control or amino acid starvation. (**d**) Alteration in cytosolic Ca2+ was measured by the Fura-2 ratio (F340/F380) in 293T cells expressing TRPML1-L15L/AA-L577L/AA at low pH (pH 4.6) external solution. Cytosolic Ca2+ alteration wasn't observed in oleic acid or fatty acid mixture (arachidonic, linoleic, linolenic, myristic, oleic, palmitic and stearic acid).

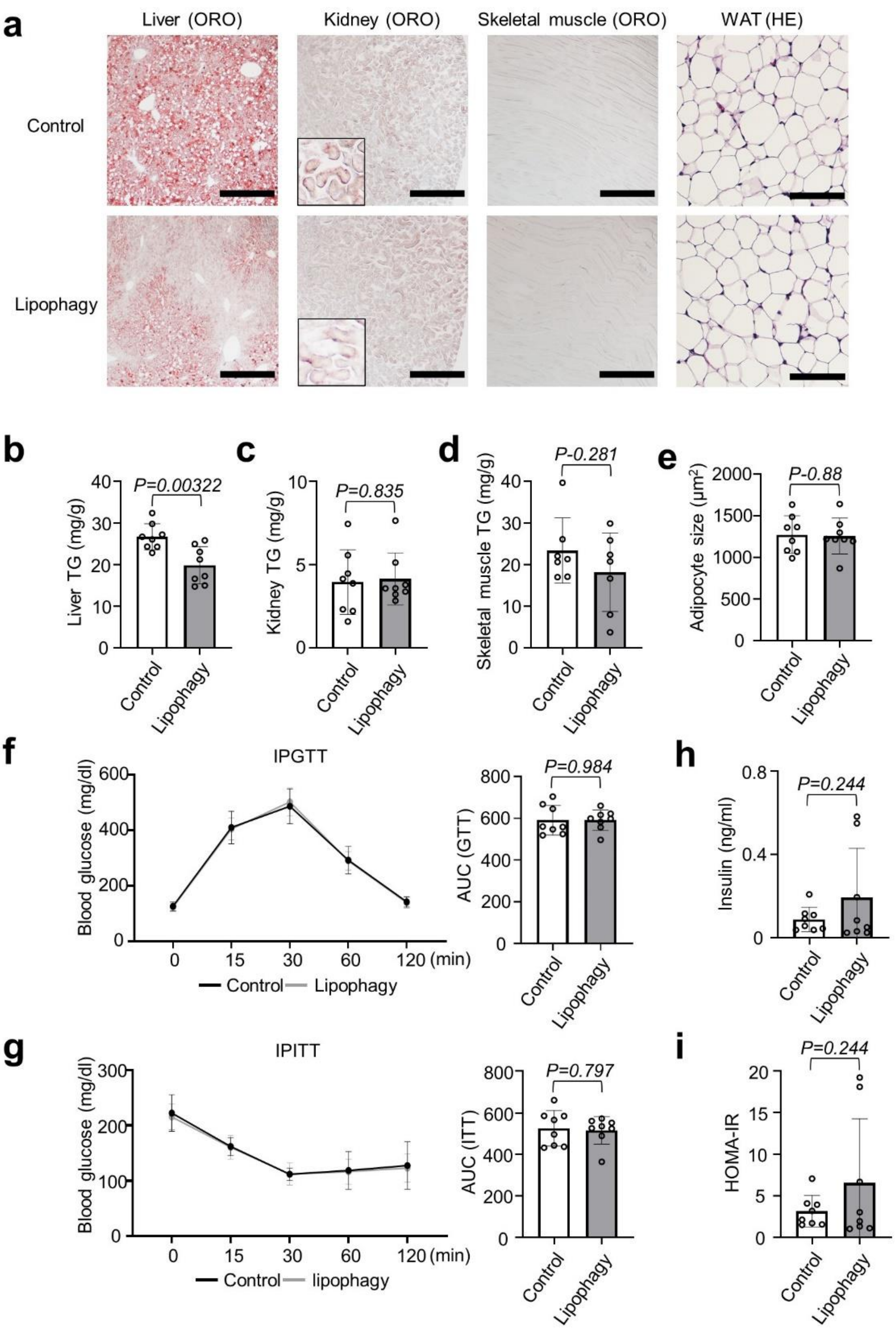


**Supplementary Figure 7. Liver lipophagy wasn't associated with ectopic fat accumulation or abnormalities glucose metabolism.** (**a**) Oil red O staining of liver, kidney and skeletal muscle sections from lipophagy-induced NASH model mice. Scale bars, 500μm. Hematoxylin-eosin (HE) staining of WAT sections. Scale bars, 100μm. (**b-d**) TG content in liver, kidney, skeletal muscle of lipophagy-induced NASH model mice. (**e**) Adipocyte size of epididymal adipose tissue. (**f-i**) Effects on the intraperitoneal glucose tolerance test (ipGTT), the intraperitoneal insulin tolerance test (ITT), the fasting serum insulin and HOMA-IR. All data are presented as mean±SD of n=8 per each group. *P* values calculated by two-sided unpaired *t*-test.
